# Paratope mapping of tilvestamab, an anti-AXL function blocking antibody, using high-throughput bacterial expression of secreted scFv-ompY fusion proteins

**DOI:** 10.1101/2024.12.12.628227

**Authors:** Eleni Christakou, Petri Kursula, David Micklem

## Abstract

Targeting AXL with a highly selective antibody presents a promising approach for inhibiting AXL and potentially improving cancer treatment. An essential step in antibody optimisation is the mapping of paratope residues to epitope residues. In this study, we identify the residues of tilvestamab, a function-blocking anti-AXL monoclonal antibody (mAb), that are essential for its binding to the extracellular domain of AXL. A single-chain variable fragment (scFv) fused to osmotically inducible protein Y (osmY) was designed to enable the secretion of soluble scFv-osmY mutants, which could be directly subjected to high-throughput biolayer interferometry (BLI) screening for binding to the AXL Ig1 domain. Each CDR residue of scFv was mutated to Ala, while additional mutations were made on the basis of predicted contribution to binding. We generated Alphafold3 predictions for the scFv(tilvestamab)-AXL Ig1 complex to gain insights into the molecular interactions of the essential residues, as determined by the experimental data. Our study reveals that tilvestamab binds to the Ig1 domain of AXL, with twelve residues on scFv (tilvestamab) contributing most to binding. Glu2 near the N terminus of AXL is essential for binding. The data give a structural view into the AXL-tilvestamab complex and allow for further optimisation of the binding interface.

## Introduction

The transmembrane protein AXL belongs to the TAM (TYRO3, AXL, MERTK) family of receptor tyrosine kinases and regulates a wide range of physiological cellular functions essential for cell survival, proliferation and migration^1^. AXL is composed of an extracellular domain, a single transmembrane domain, and a cytosolic tyrosine kinase domain. The N-terminal extracellular domain interacts with extracellular ligands to initiate signal transduction. This ectodomain consists of two immunoglobulin-like (Ig like) domains and two fibronectin type III (FNIII) domains. A single transmembrane domain is followed by an intracellular domain harboring tyrosine residues conserved among the TAM receptors, as well as the kinase domain. The Ig domains within the ectodomain of AXL interact with its primary ligand, growth arrest-specific protein 6 (GAS6), leading to receptor dimerization, autophosphorylation of the tyrosine residues and activation of the intracellular signaling cascades^2^.

Axl expression is upregulated in numerous human malignancies, such as leukemia, breast cancer, prostate cancer^3^, kidney cancer^4^, melanoma^5^, glioblastoma^6^, and non-small lung cancer^7^, and it is associated with cancer progression, therapy resistance and poor prognosis^8–11^. Given its role in cancer development, AXL has emerged as a promising prognostic marker and therapeutic target. AXL-targeted therapies with small molecule inhibitors, which target the kinase domain in either its active or inactive conformation, have shown promising results and have progressed into various phases of clinical investigation^12–14^. Antibody therapies present an alternative approach to small-molecule inhibitors with higher selectivity thereby minimizing off-target toxicity. High affinity anti-AXL monoclonal antibodies (mAb) exert tumor inhibition either as a single agent or in combination with known chemotherapeutic drugs^15^.

Tilvestamab (BGB149) is a function-blocking, humanized and highly selective anti AXL-mAb, developed by BerGenBio^16^. In this study, we investigated the essential amino acids that are involved in the interaction of tilvestamab with its target antigen, AXL. The variable V_H_ and V_L_ domains of antibodies each contain three hypervariable loops, known as complementarity-determining regions (CDR), which pack together at the surface of the antibody to form the paratope that binds to the antigen epitope. To map the paratope of tilvestamab, we generated a panel of bacterially expressed single-chain fragment variables (scFvs) carrying tilvestamab CDRs with various single point mutations and screened them for their ability to bind the Ig1 domain of human Axl. The set of point mutations included mutation of each CDR residue to alanine and additional mutations of key residues (as defined in ^17^) to a wider set of substitutions. To enable rapid functional screening of the antibody fragments, the scFvs were fused to the secreted bacterial osmotically inducible protein Y (osmY) allowing testing of protein directly in the culture supernatant^18^. Finally, we demonstrate how the paratope, defined by the experimental data, aligns to the AlphaFold structural prediction. The updated AlphaFold3 exhibits higher accuracy in modeling antibody-antigen complexes compared to its predecessor, AlphaFold2^19^. The 3D structural model predicted using AlphaFold3 reveals how the key amino acids identified in the screen interact at the molecular level, highlighting the specific interactions that drive the formation of the antibody-antigen complex. Taken together, the results can assist the design of second-generation therapeutic antibodies with improved properties.

## Materials and Methods

### Design of plasmid encoding scFv-osmY fusion protein

The sequence of tilvestamab was reconfigured into a shorter scFv antibody format, incorporating the variable domains of the heavy (V_H_) and the light (V_L_) chains, joined together by a short flexible peptide linker AKTTPPKLEEGEFSEARV (adapted from the vector pOPE101, Progen, Genbank, accession Y14585). The scFv was connected by a short linker RADAAPTVSAA to the gene coding for the bacterial osmotically inducible protein Y (osmY). This was followed by a Myc-tag (GSEQKLISEEDL), facilitating detection of the fusion protein by the mouse monoclonal antibody mab Myc1-9E10, and a His_6_-tag. The sequence was synthesized and cloned into the bacterial T7-driven expression vector pET-22b(+) by Genscript. All plasmid DNA sequencing was carried out at Eurofins Genomics.

All protein sequences and primers used are listed in the **Supplementary Material**. The scFv-osmY fusion protein expression system allows for high-throughput, low-effort expression of scFv secreted directly into the medium (not just the periplasmic space) ^18^.

### Mutagenesis

scFvs were diversified by mutagenesis as described in the Q5 site-directed mutagenesis kit (New England Biolabs). Mixtures of mutagenic primers were prepared at 1 µM for each primer. Plasmid template (pET-22b-osmY-scFv) was diluted to 5 ng/µl in water, and then to 50 pg/µl in 2x Q5 Mastermix. Then, in a PCR plate, 4 µl of Mastermix and 4 µl of the appropriate primer mix were distributed to each well. Reactions were run in the following thermal cycler program: 98 °C for 30 s; 24 cycles [98 °C for 10 s; anneal for 10 s; 72 °C for 210 s]; 4 °C hold. The annealing temperature was chosen according to the primer pair, using a gradient on the thermocycler when appropriate. Products were treated with the KLD mix to degrade the parental vector, add 5’ phosphates to the DNA products, and ligate the linear DNA into circular plasmids. A master mix was made by adding 1/5 vol KLD enzyme mix to the 2x KLD reaction buffer and distributed as 1 µl aliquots. 0.7 µl of each PCR product diluted 1+3 in water was added and mixed. After incubation for at least 5 min at room temperature, 5 µl of the KLD reaction was transformed into chemically competent T7 express *E. coli* cells via heat shock, plated on LB + carbenicillin plates and incubated at 37 °C overnight.

### Bacterial expression of tilvestamab scFv fragments

Autoinduction is a method to induce protein expression from T7-lacO-regulated promoters without requiring the addition of the expensive lactose-analog, isopropyl β-D-thiogalactopyranoside (IPTG). It is based on the observation that *E. coli* will preferentially use glucose and glycerol as an energy source even in the presence of lactose, and that these carbon-sources will maintain lacO in an inactive form. Only when the glucose and glycerol have been used up (and when the culture is at a high density) will the lactose allow T7-mediated expression^20^.

Autoinduction was accomplished by inoculating single colonies of a bacterial T7 expression strain carrying an scFv-ompY fusion protein construct (pET-22b-osmY-scFv or mutated variants) into 1 ml of Autoinduction medium (LB Broth supplemented with 0.5% glycerol, 0.05% glucose, 0.2% lactose, 50 µM Fe(III)(NO_3_)_3,_ 20 µM CaCl_2,_ 2 µM NiCl_2,_ 2 µM CoCl_2,_ 2 µM boric acid, 25 mM Na_2_HPO_4,_ 25 mM KH_2_PO_4,_ 25 mM (NH_4_)_2_SO_4_ and 100 µg/ml carbenicillin) and grown with vigorous shaking at 37 °C for at least 24 h. Bacteria were pelleted by centrifugation at top speed in a benchtop microcentrifuge. Pellets were retained for DNA preparation and sequencing, and the scFv-ompY fusion protein-containing supernatant was used directly in Western blotting and Octet binding experiments.

### Detection of scFv-ompY proteins by Western blotting

For each mutant, bacteria were pelleted and 10-40 µl bacterial supernatant was run on an SDS-PAGE MiniPROTEAN TGX stain-free 4-20% gel (BioRad) and blotted onto a polyvinylidene difluoride membrane (GE Healthcare Life Sciences). The membranes were blocked with 10 ml of 5% (w/v) skimmed milk in Tris-buffered saline/Tween-20 (TBS-T; 25 mM Tris-HCl, 1 mM NaCl, 0.1% Tween-20) followed by overnight incubation at 4 °C in 5 ml of incubation buffer (TBS-T with 5% skimmed milk) containing biotinylated anti-Myc primary antibody (Millipore 16-170) in 1/1000 dilution. The membranes were then washed three times with PBS-T (0.01% Tween20 in PBS) and incubated with the secondary reagents HRP-Streptavidin (BD Pharmingen 51-75477E, further diluted 1/1000) and HRP-StrepTactin (BioRad Cat#1610380, diluted 1/10,000) in 5 ml of incubation buffer for 20 min-4 h at RT. Finally, after three washes with PBS-T, protein bands were visualised with enhanced chemiluminescence (Pierce^TM^ ECL Substrate, 32106) and imaged using a ChemiDoc^TM^ XRS+ imager with Image Lab software (Biorad Hercules, CA).

### Biolayer interferometry screening of scfv-osmY supernatants

Mutated scFv-osmY were screened for binding to the Ig1-domain of huAXL by biolayer interferometry (BLI) on an Octet RED96 instrument (Pall ForteBio). Reagents and samples were dispensed into 96-well polypropylene, black flat-bottom microplates (Greiner). Dip and Read™ Ni-NTA coated biosensor tips (ForteBio) were used to capture the His-tagged scFv-osmY proteins directly from the undiluted bacterial supernatants. The method used consisted of the following steps: (1) equilibration of tips in kinetic buffer (0.02% Tween 20, 0.05% sodium azide, 0.1% BSA in PBS) to acquire baseline (120 s); (2) loading of scFv-ompY by transferring the biosensors into wells containing undiluted supernatant (600 s); (3) rinsing of tips (5 s) to remove non-specific proteins and baseline acquisition in kinetic buffer (120 s); (4) association of AXL Ig1 domain Fc (5.6 μg/ml in kinetic buffer); (5) dissociation by dipping the biosensors into wells containing kinetic buffer; (6) regeneration of biosensors by immersion in 10 mM glycine (pH 1.5); and neutralisation in kinetic buffer (3 * 5 s cycles); (7) recharging of biosensors with 10 mM NiCl_2_ (60 s).

Binding was analysed by isolating the Association/Dissociation phases and aligning them on the y-axis by subtraction of the baseline acquired in step 4. Individual BLI traces were scored by eye to determine whether the scFv-ompY being tested bound apparently normally to AXL-Ig1-Fc (Normal), failed to bind at all (None), or showed an intermediate effect. “Fast” and “Very Fast” describe scFv-ompY strong initial binding during the association phase, followed by rapid or very rapid loss of signal (compared to the parental scFv-ompY). In addition, some scFv-ompY were scored as “Weak” or “Weak Fast”; these showed detectable but weak initial binding followed by slow or rapid dissociation. Examples of binding curves are shown in Fig.1.

**Figure 1.**
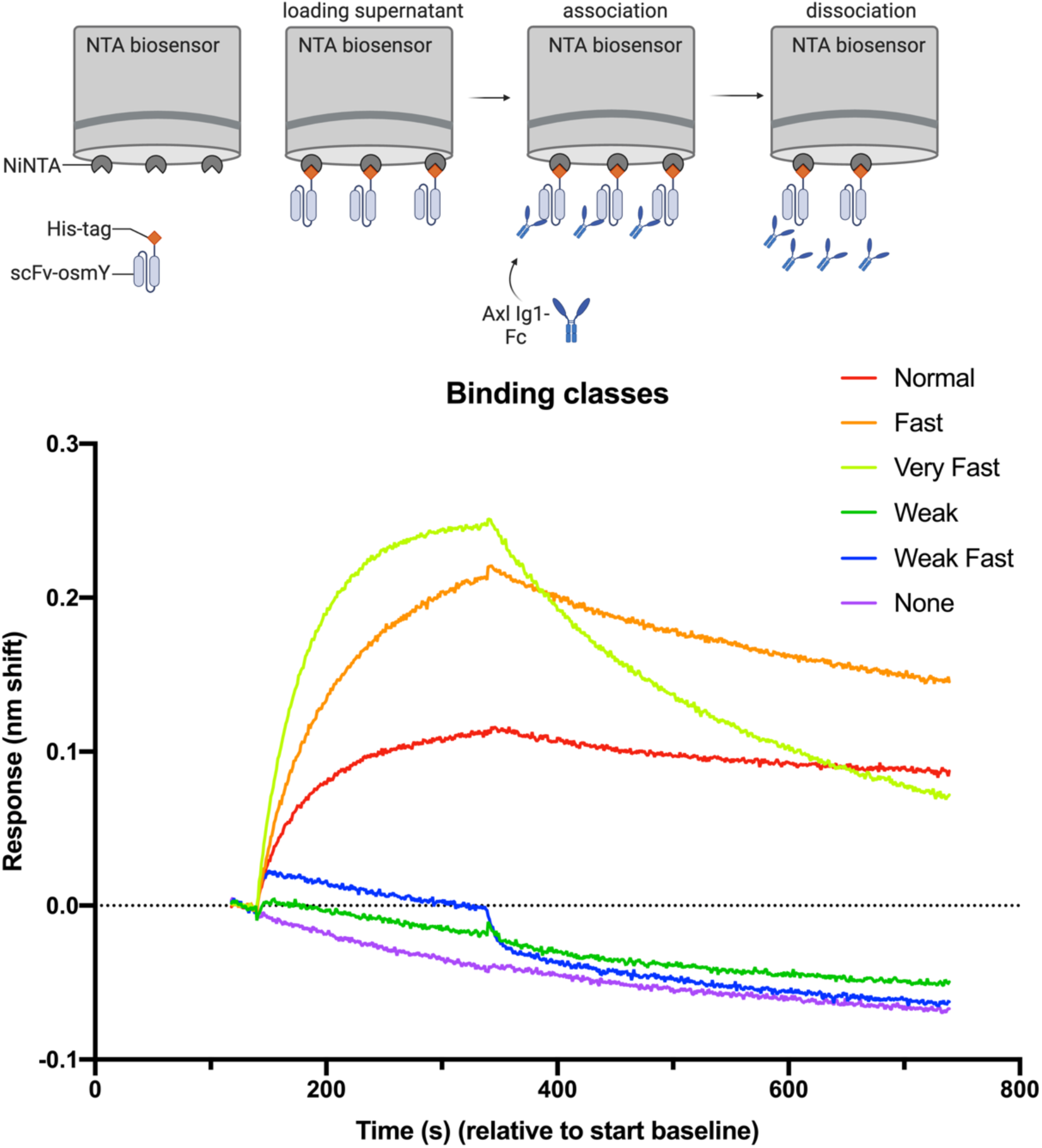
Schematic representation of the BLI strategy. Top: NTA biosensors were used to capture scFv-osmY mutants and test their binding to AXL Ig1-Fc. Bottom: the representative binding sensorgram shows different examples of binding curves from the BLI experiments and how the binding was classified as normal, fast, very fast, weak, weak fast and no binding.

The strength of binding was not obviously related to the expression level of the scFv-ompY as determined by Western blotting. To visualize the binding information in a more condensed form, a “Severity score” was assigned to each binding conclusion to indicate how severely binding was affected. According to the scoring system, 0, 1, 2, or 3 points were assigned to normal, fast, very fast, and weak binding, and the higher scores 4 and 5 indicated very weak or weak fast, or no binding. Substitutions that were scored “normal” or “fast” were considered tolerated, while all other severities were considered not tolerated.

### Protein structure prediction and interpretation

An AlphaFold3 model of the AXL-Ig1 – scFv complex was generated at the AlphaFold3 server^19^. For comparison, an AlphaFold2^21^ model was made using the AlphaFold2 Colab^22^. Visualisation and analysis of the structure were carried out in PyMOL^23^.

## Results and Discussion

The identification of essential residues of tilvestamab for binding to the AXL Ig1 domain involved the following steps (illustrated in **Fig.2**). First, a scFv-osmY antibody-like protein was designed, carrying the variable regions of tilvestamab. Next, all residues within the CDR loops were mutated to alanine, while additional mutations were made on selected residues guided by the work of Robin et al. predicting certain residues to contribute most to the binding energy^17^. Mutants were sequenced and tested individually for binding to AXL Ig1-Fc protein by BLI. After experimentally scoring the mutations, an AlphaFold3 model of the complex was generated for structural insights.

**Figure 2.**
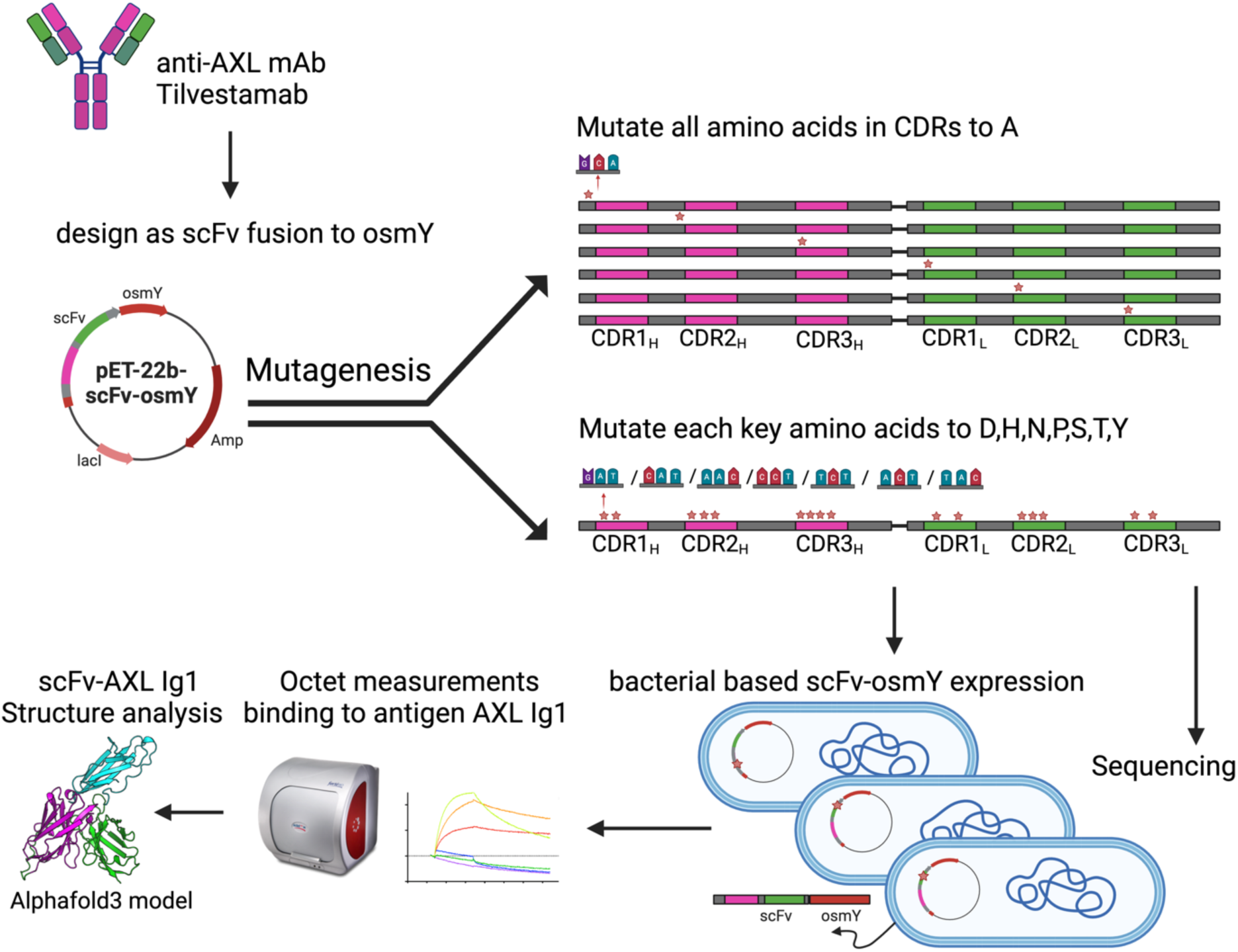
Overview of the experimental strategy for the identification of essential residues of tilvestamab for binding to AXL. Site-directed mutagenesis was used to alter each CDR residue to alanine, while additional mutations were also made on selected residues. BLI was used to test each mutant binding to the Ig1 domain of AXL. Finally, the experimental data were linked to AlphaFold3 prediction of the complex scFv-Ig1 AXL.

### Tilvestamab expressed as a single-chain antibody fragment fused to ompY binds the AXL Ig1 domain

The complementarity determining regions of tilvestamab were reconfigured as a single-chain antibody fragment (scFv) fused to the bacterial protein osmY carrying Myc and His6 tags for high-throughput production of antibodies in *E. coli* (parental plasmid map, Supplementary Fig.1). The protein produced from this vector is referred to as “parental”. Following expression in a T7-expressing strain of *E. coli,* the approximately 52-kDa protein could be readily detected in the culture supernatant by Western blotting with an anti-Myc antibody (**Fig. 3A**).

**Figure 3.**
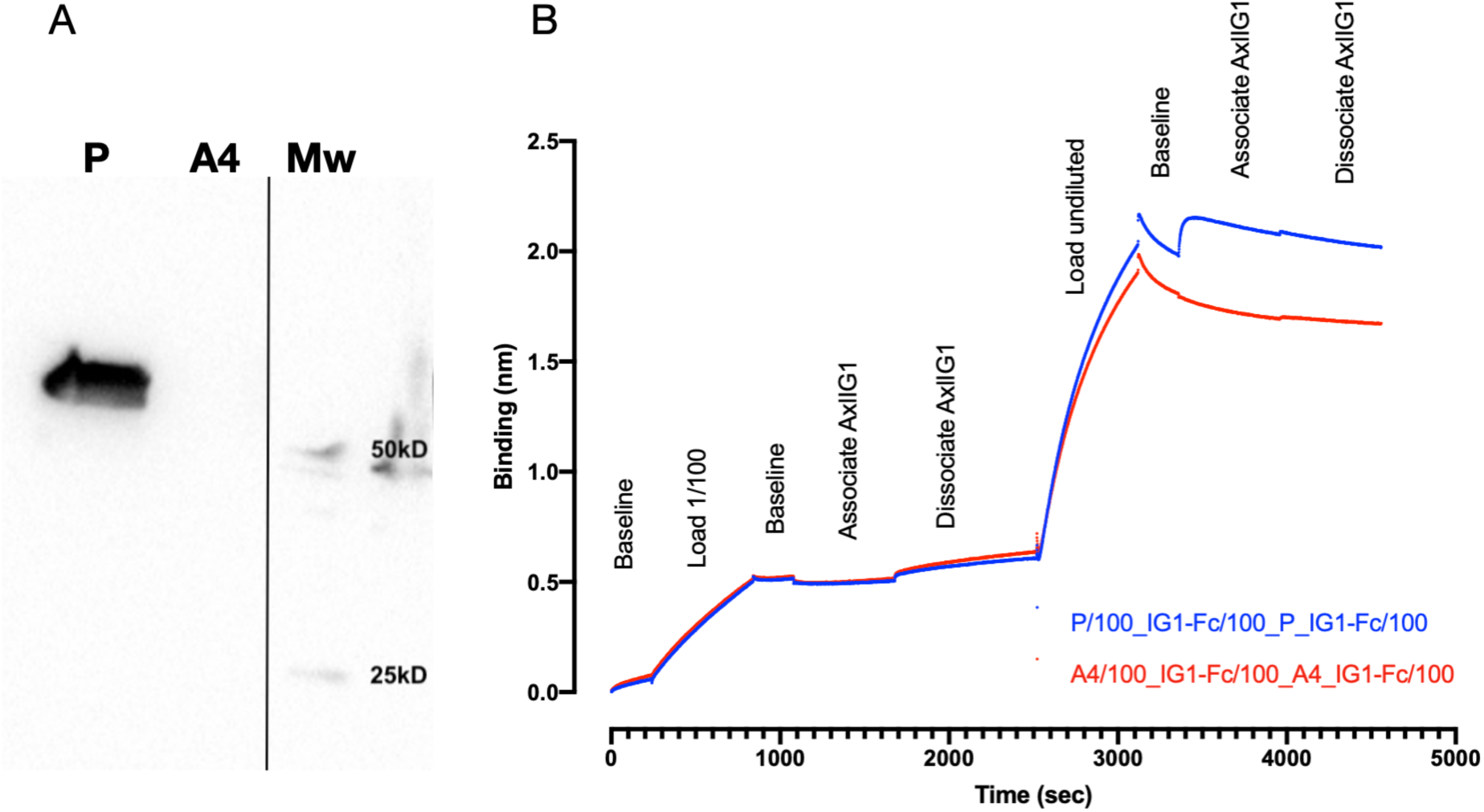
Parental scFv-ompY expressed in bacteria binds to AXL Ig1 domain. **(A)** Western blot with anti-Myc antibody detects a strong band in supernatant from bacterial cells expressing Parental scFv-ompY (P) but not from control supernatant from cells expressing a frame-shifted mutation (A4). Mw: Molecular weight markers. **(B)** Biolayer interferometry analysis on an Octet RED96 demonstrates binding of P (but not A4) from undiluted culture supernatant to human AXL IG1-domain-Fc fusion protein (“Associate AXLIG1”). Supernatant used at 1/100 (first 2500s) did not give a detectable response.

The His6-tag on the scFv-ompY fusion proteins allows them to associate with NiNTA-coated tips for BLI experiments (**Fig. 3B**). Tips loaded with undiluted culture medium from cells expressing the parental tilvestamab-like scFv-ompY bound strongly to the AXL Ig1 domain fused to human Fc (blue, t=3300 s onwards, **Fig. 3B**). Control supernatant (frameshift mutated scFv-ompY) demonstrated loading of unspecific proteins onto the tips (t=2500-3100 s), but no association with AXL Ig1 Fc (red, t=3300 s onwards). No specific interaction could be seen when culture supernatant was used diluted 1/100 (t=0-2500 s). As expected, parental scFv was unable to bind rat AXL Fc (not shown). Thus, the parental scFv-ompY fusion protein based on the antibody recognition sites of tilvestamab retained the ability to specifically bind the human AXL Ig1 domain.

### Design of a panel of mutated scFv-ompY variants

Robin et al. computationally identified the residues within antibody CDRs that are most likely to contribute energetically to antigen binding^17^. Using the worksheet that they published, we identified residues within the CDRs that are likely to contribute to binding (**Fig. 4**). These residues were mutated using degenerate oligonucleotides with the appropriate codon replaced by NMT (AGCorT, AorC, T), which encodes an amino acid from the set alanine (A), aspartate (D), histidine (H), arginine (N), proline (P), serine (S), threonine (T) and tyrosine (Y), representing a broad subset of amino acid classes including charged (DH), polar (DHNSTY), hydrophobic (AHTY), hydrophilic (DNPS), aromatic (Y), tiny (SA), and small (ADNSP). In addition, a small number of mutations to leucine were generated. All other residues in the CDRs were individually mutated to alanine. The set of primers used for site-directed mutagenesis is listed in **Supplemental Table 1 and 2**.

**Figure 4.**
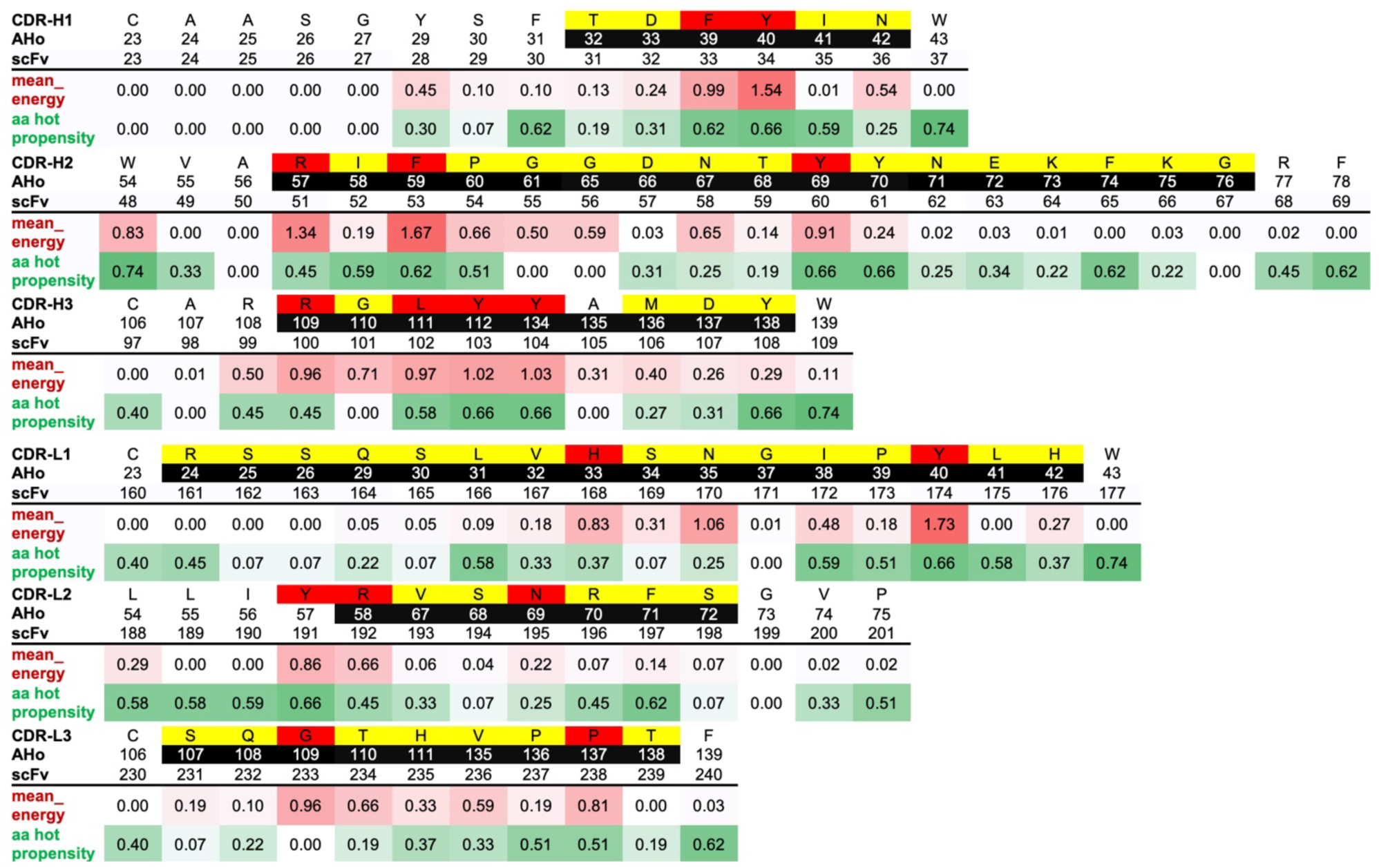
Mutations of Heavy Chain CDRs (CDR-H) and Light Chain CDRs (CDR-L) from tilvestamab. Tilvestamab seq: The sequence of each CDR and surrounding framework-region sequence. AHo: the numbering of the amino acid according to the numbering scheme of Honegger and Plückthun^24^. CDR residues as defined by Kabat are shown as white text on black shading. scFv: numbering of the amino acid within the scFv-ompY fusion protein. mean_energy: the average binding free energy contributed by each site^17^. aa hot propensity: a measure of how often an amino acid makes an energetically favorable bond when it is in a hot-spot location in the sequence adapted from Robin et al^17^. Amino acids targeted for mutation to alanine are shaded yellow. Amino acids targeted for mutation to ADHNPSTY are shaded red.

### Scoring scFv-osmY variants

scFv-ompY variants were expressed in *E. coli* and analysed using BLI. Each scFv-ompY variant protein was first loaded onto the NiNTA sensor tips, and the association and dissociation of human AXL Ig1-Fc protein to the immobilised scFv-ompY was then followed, leading to assignment of a Severity Score for each mutation. In **Fig. 5A-F**, the results for each scFv-ompY variant are plotted. Overall, all six CDR loops of the tilvestamab scFv were important for the binding interaction, while for both the heavy and light chains, CDR2 appeared to tolerate the most mutations.

**Figure 5.**
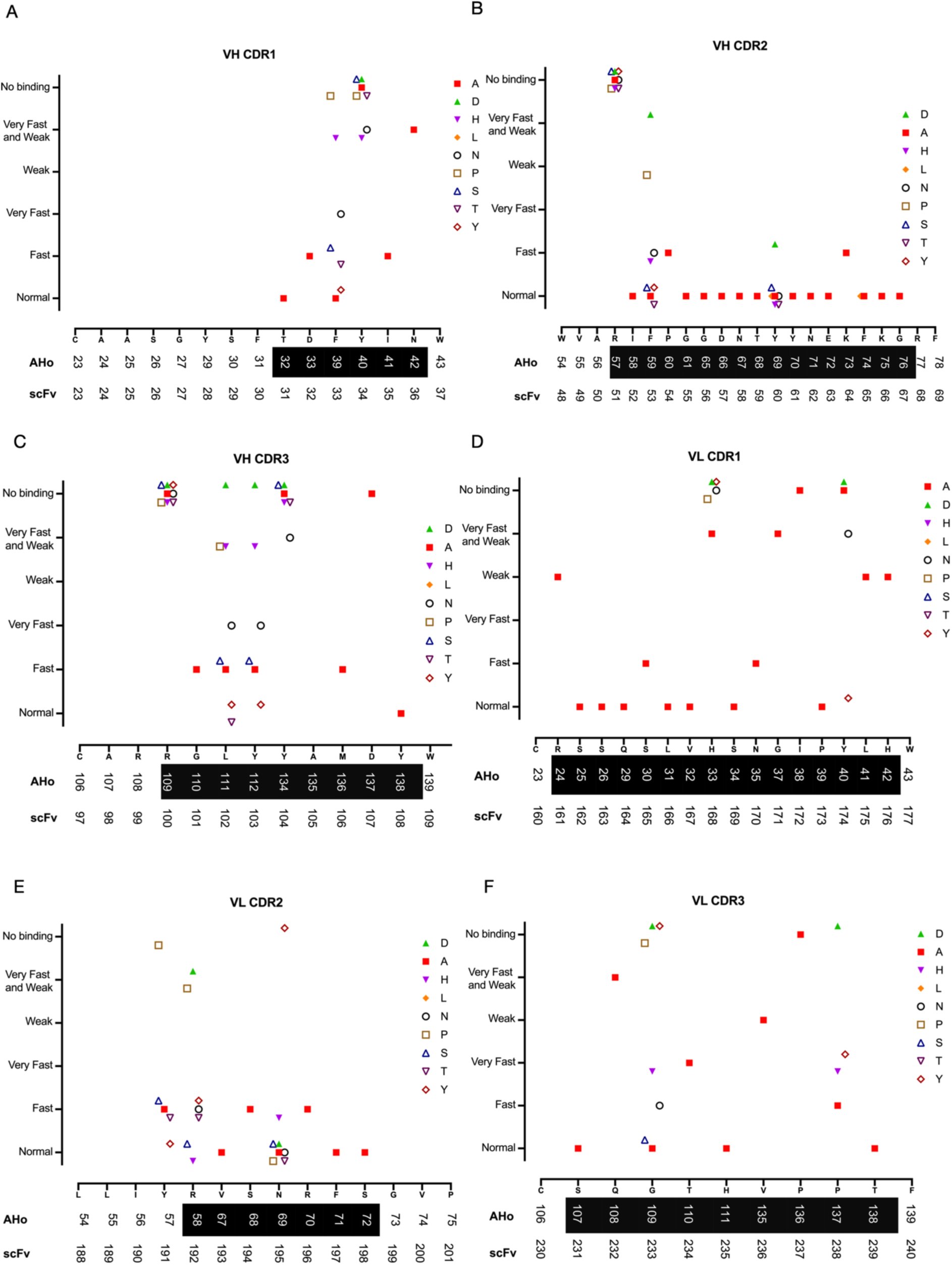
Effect of mutations binding to AXL Ig1-Fc for all six CDR loops of tilvestamab scFv **(A)** heavy chain CDR1, **(B)** heavy chain CDR2, **(C)** heavy chain CDR3, **(D)** light chain CDR1, **(E)** light chain CDR2 and **(F)** light chain CDR3. The results are plotted with severity score on the y-axis vs the original sequence on the x-axis. AHo numbering, the location of each CDR and the numbering of the amino acid within the scFv are also indicated. AHo: the numbering of the amino acid according to the numbering scheme of Honegger and Plückthun^24^. CDR residues as defined by Kabat are shown as white text on black shading. scFv: numbering of the amino acid within the scFv-ompY fusion protein.

The results shown in **Fig. 5A** suggest that the Tyr at AHo 40 (scFv Y34) and the Asn at AHo 42 (N36) are essential for binding. AHo 39 (scFv F33) tolerates the conservative substitution to Y, and the small amino acids A, S and T but not P, H or N. T31, D32, I35 all tolerate substitution to Ala suggesting that they do not contribute greatly to binding energy.

The results shown in **Fig. 5B** suggest that the Arg at AHo 57 (scFv R51) is essential for binding. Substitution to Ala has only a minor effect for all other amino acids in CDR2. AHo 59 (F53) tolerates most tested substitutions except for Pro and the negatively charged Asp.

The results shown in **Fig. 5C** suggest that several amino acids in the heavy-chain CDR3 are important for binding: the Arg at AHo 109 (scFv R100), the Tyr at AHo 134 (scFv Y104) and the Asp at Aho 137 (scFv D107) do not tolerate substitution to any of the tested amino acids. Substitution to Ala has only a minor effect for AHo 110 (scFv G101), AHo 136 (scFv M106) and AHo 138 (scFv Y108). AHo 111 (scFv L102) and AHo 112 (scFv Y103) tolerate only some substitutions. In both cases, substitutions to Ala, Ser, or Tyr are tolerated, while substitution to Arg, His, or Asp causes progressively worse binding. scFv L102 was also tested for substitution to Thr (tolerated) and Pro (not tolerated).

In the light chain CDR1 (**Fig.5D**) AHo residues 24, 33, 37, 38, 40, 41, 42 (scFv R161, H168, G171, I172, Y174, L175 and H176) are all essential for strong binding and do not tolerate substitution to Ala or any other amino acid tested.

In the light chain CDR2 (**Fig.5E**) all CDR residues tolerate substitution to Ala (not tested for AHo 58, scFv R192). Most other substitutions are well tolerated apart from AHo 57 (scFv Y191) mutation to Pro, AHo 58 (scFv R192) mutation to Pro or Asp, and AHo 69 (scFv N195) mutation to Tyr.

Finally, in the light chain CDR3 (**Fig.5F**), four CDR residues do not tolerate substitution to Ala (AHo 108, 110, 135, 136; scFv Q232, T234, V236, and P237). In addition, AHo 109 and AHo 137 (G233 and P137) do not tolerate substitution to Asp, Tyr, His, or Pro (G233 only), while tolerating Ala.

### Structural interpretation of the experimental data based on AlphaFold3 predictions

Based on the experimental alanine scanning data, the paratope residues that contributed most to binding – those that either abolished all binding or resulted in very weak binding with quick dissociation – are summarized in **Table 1**. Where other amino acids were tested at these locations, they were also not tolerated (**Fig. 5**). To support the understanding of these essential interactions within the complex, we utilized AI-based prediction using the latest AlphaFold3 (**Fig. 6A,B**). The model and the specific molecular interactions that help stabilize the complex are shown in **Fig. 6**. Contact residues of the V_H_ CDR loops can be seen buried within the interacting interface, participating in hydrogen bonds and van der Waals interactions with the AXL Ig1 domain.

**Figure 6.**
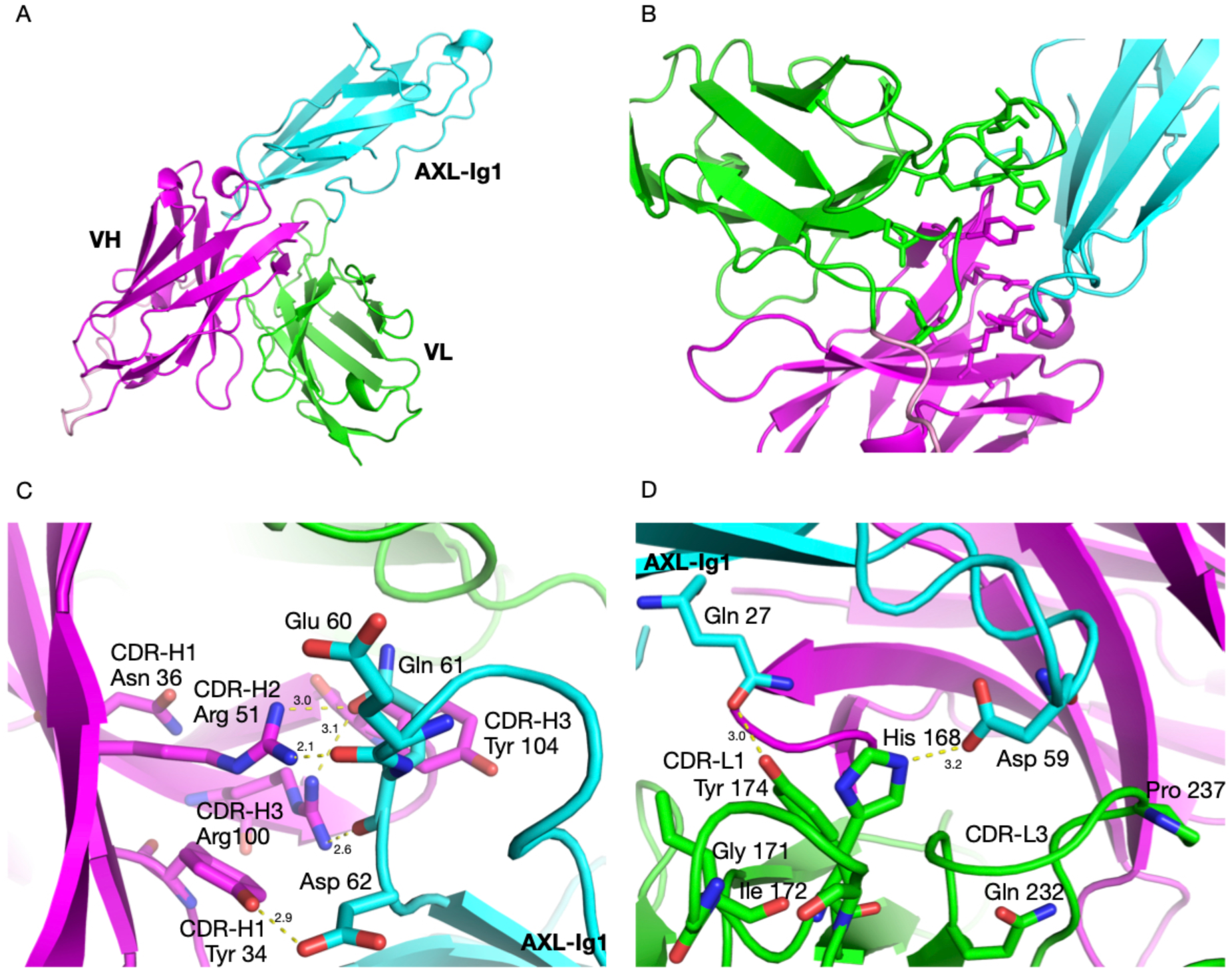
AlphaFold 3 structure prediction and analysis. **(A)** Overall view of AXL Ig1-scFv complex. AXL Ig1 domain is indicated in cyan. The VH and VL of tilvestamab are colored magenta and green, respectively **(B)** Tilvestamab paratope with the side chains of the 12 key amino acids as identified by the alanine-scanning experimental data. (C,D) Key amino acids interactions between AXL Ig1 and VH domain **(C)** and VL domain **(D)**. Oxygen and nitrogen atoms of the side chains are colored red and blue, respectively, and hydrogen bonds between AXL Ig1 and scFv are indicated with yellow dotted lines. Numbering of AXL Ig1 amino acids according to **Supplemental Fig.2**.

**Table 1.**
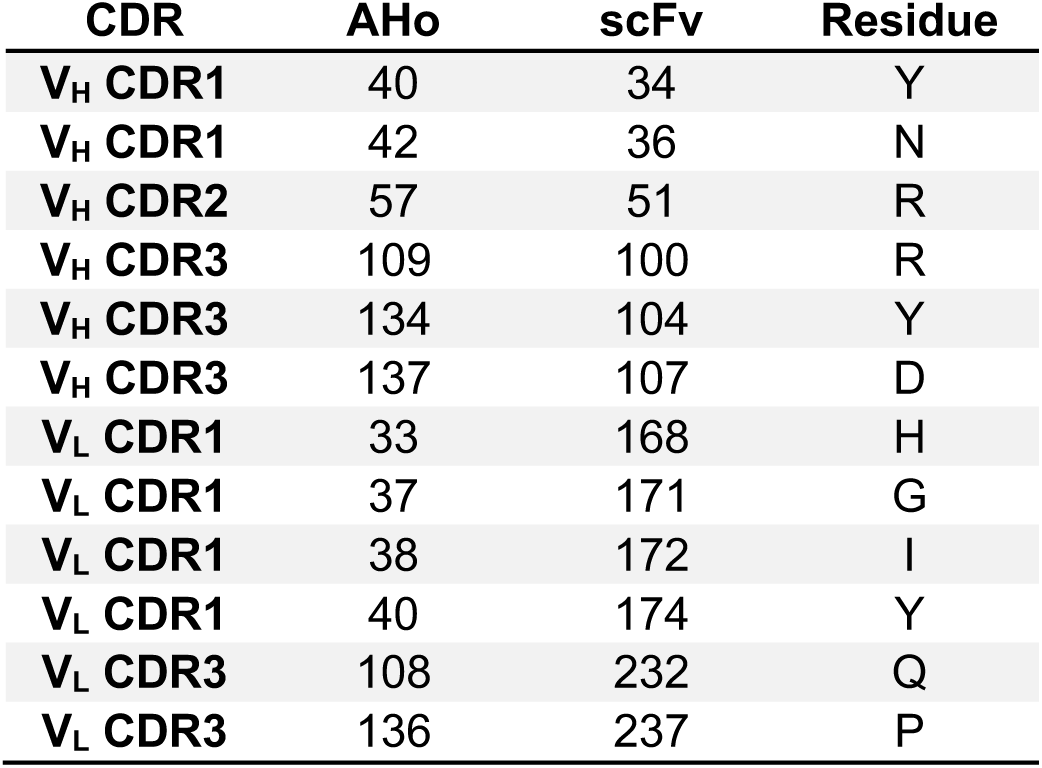
Key amino acids of tilvestamab contributing to binding of AXL Ig1 domain. AHo: the numbering of the amino acid according to the numbering scheme of Honegger and Plückthun^24^. scFv: numbering of the amino acid within the scFv-ompY fusion protein.

The hydroxyl group in the side chain of Tyr34 forms a hydrogen bond with the carboxylate group in the sidechain Asp62 of AXL Ig1 (numbering of AXL Ig1 according to **Supplementary** Fig.2). The guanidinium group, in the vicinity of Tyr34, of Arg51 forms hydrogen bonds with the backbone carbonyl group of Glu60 and the side-chain carbonyl oxygen of Gln61. Moreover, in CDR3, Arg100 nearby forms two hydrogen bonds with the backbone and side-chain carbonyl oxygens of Gln61, while the side chains of Tyr104 and Asp107 contribute to van der Waals interactions at the interface that would weaken upon mutation to Ala (**Fig. 6C**).

Direct interactions of AXL Ig1 are less prevalent with residues of the V_L_ CDR loops. The nitrogen in the histidine imidazole group of His168 forms a hydrogen bond with the carboxyl group in Asp59 of AXL Ig1 (**Fig. 6D**). Another hydrogen bond is formed between the hydroxyl group in the aromatic ring of Tyr174 and the side chain amide of Gln27, while the hydrophobic side chain of Ile172 residue being near the also hydrophobic Ile65 of AXL Ig1 can participate in van der Waals interactions. The remaining residues listed, Gly171, Gln232, and Pro237, do not tolerate mutations to Ala, as such changes could disrupt CDR loop flexibility and conformation.

Sites that tolerated mutation to alanine but showed variable tolerance for other substitutions are listed in **Table 2**. Specifically, when looking at the AlphaFold3 model of scFv-AXL Ig1, Phe33 has close interactions with AXL through its aromatic side chain. Mutations to Ala or Tyr at that side had no effect (**Fig. 5A**), indicating a hydrophobic and/or aromatic residue at this position is important. The F33H and F33N mutants make the side chain – and the surface – more hydrophilic, while Pro at this site would affect the conformation of the entire CDR1 loop. Phe53 nearby also interacts with AXL via its side chain. Most mutations are tolerated, except for substitution to Pro and the acidic Asp; these substitutions could affect the local conformation and surface charge, respectively. Lys102 and Tyr103 in V_H_ CDR3 both make van der Waals contacts to AXL, and Tyr103 also makes a hydrogen bond to the carbonyl of Pro4 in AXL. In the experiment, most mutations in Lys102 and Tyr103 affected binding, which can be explained by the loss of these interactions. The most significant impact being the mutation to Asp, introducing an unfavourable negative charge into the binding site.

**Table 2.** Additional amino acids of tilvestamab contributing to binding of AXL Ig1 domain. The sites listed, tolerate mutation to alanine, but not to the above substitutions. AHo: the numbering of the amino acid according to the numbering scheme of Honegger and Plückthun^24^. scFv: numbering of the amino acid within the scFv-ompY fusion protein.

| CDR | AHo | scFv | Residue | Substitutions |
| --- | --- | --- | --- | --- |
| V<sub>H</sub> CDR1 | 39 | 33 | F | H, N, P |
| V<sub>H</sub> CDR2 | 59 | 53 | F | D, P |
| V<sub>H</sub> CDR3 | 111 | 102 | L | D, H, N, P |
| V<sub>H</sub> CDR3 | 112 | 103 | Y | D, H, N |
| V<sub>L</sub> CDR2 | 57 | 191 | Y | P |
| V<sub>L</sub> CDR2 | 58 | 192 | R | D, P |
| V<sub>L</sub> CDR2 | 69 | 195 | N | Y |
| V<sub>L</sub> CDR3 | 109 | 233 | G | D, H, P, Y |
| V<sub>L</sub> CDR3 | 137 | 238 | P | D, H, Y |

Tyr191, Arg192, and Asn195 in V_L_ CDR2 appear to form a structurally critical region for the complex. The most drastic effects arise when Tyr191 or Arg192 are mutated to Pro and Asp, indicating importance for the correct conformation of CDR2 and lack of tolerance for a negative charge at the paratope. Results in the Supplementary Material show that tilvestamab binds to the Ig1 domain of human AXL and rhesus monkey AXL, but not mouse AXL protein (**Supplementary Figure 3**), and a single glutamate, Glu2, close to the N terminus of the mature AXL protein, is essential for binding (**Supplementary Figure 4**). scFv-AXL Ig1 complex modelled by AlphaFold3 agrees with Glu2 of AXL being important for binding tilvestamab, as Tyr191 and Ser198 form hydrogen bonds with Glu2, and the hydrophobic Phe197 additionally stabilizes the protein interface (**Fig. 7**). AlphaFold2 models AXL Ig1 in a different orientation, with Glu2 not interacting with tilvestamab, whereas AlphaFold3 apparently improves the accuracy of the protein complex, aligning with the experimental observations and the importance of Glu2 (**Fig. 7**).

**Figure 7.**
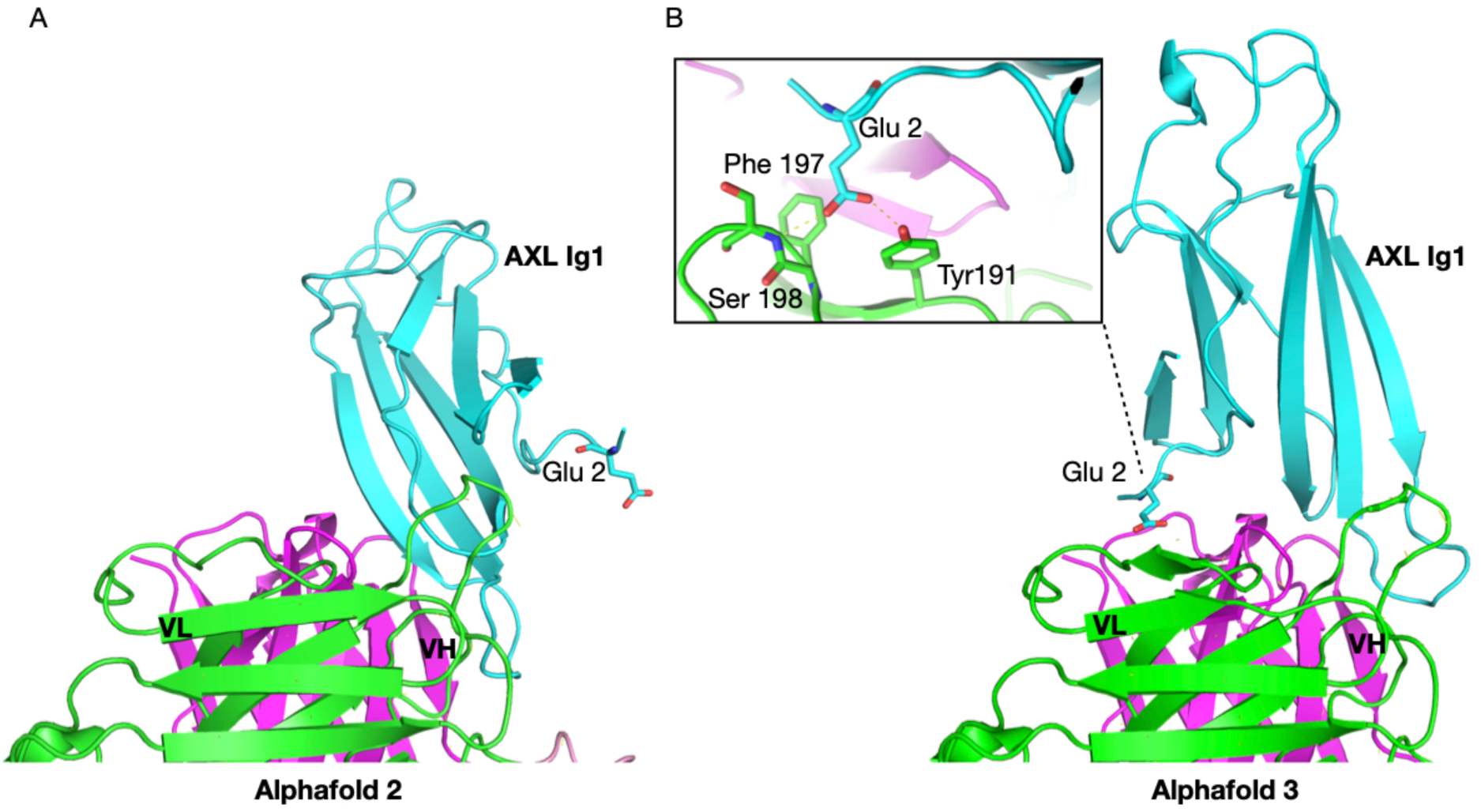
AlphaFold 2 and AlphaFold 3 comparison predicting scFv-AXL Ig1 complex from their amino acid sequence. **(A)** AlphaFold 2 structural model shows the N-terminus of AXL not interacting with scFv tilvestamab, whereas **(B)** AlphaFold 3 provides a more accurate structure prediction, modeling Glu2 interacting with residues of tilvestamab. This prediction aligns with the experimental work. Numbering of AXL Ig1 amino acids according to **Supplemental Fig.2**. The protein coloring is consistent with Fig. 6.

Lastly, in the V_L_ CDR3 loop, Gly233, and Pro238 appear structurally critical, but neither makes direct contact with AXL. These are the most flexible and most rigid amino acids. With the hydrogen atom side chain of Gly233 pointing towards the heavy chain, substitutions with smaller amino acids S and A would be tolerated but larger or charged residues such as D, H, P, or Y would be more disruptive in that position as reflected in the experimental binding data. In addition, Gly233 carbonyl group forms a hydrogen bond with Gln61 of AXL, and the geometry of this bond would be affected by mutations. Mutations at Pro238, located in the end of the CDR3, affect the most rigid amino acid, and all mutations influence binding likely due to changes in conformation of the CDR3 loop.

Tilvestamab is known to block AXL signalling ^25,26^, while it reduced GAS6 binding to AXL only by ∼50% ^16^. This suggests that tilvestamab does not simply block GAS6 binding to AXL but affects the formation of a signalling-competent AXL dimer upon GAS6 interaction. To get an insight into this subject, we superposed the AlphaFold3 prediction of the AXL Ig1 – tilvestamab scFv complex on the crystal structure of the complex between AXL and GAS6 ^27^. The superposition (**Fig. 8**) shows that the binding sites for tilvestamab and GAS6 lie close on the surface of AXL Ig1, with a small overlap. The N terminus of AXL, with Glu2, does not interact with GAS6 in the crystal structure. These observations suggest that tilvestamab may reduce the binding affinity of GAS6 and affect the conformation and oligomeric state of the protein complex, such that a signalling-competent AXL receptor dimer cannot form.

**Figure 8.**
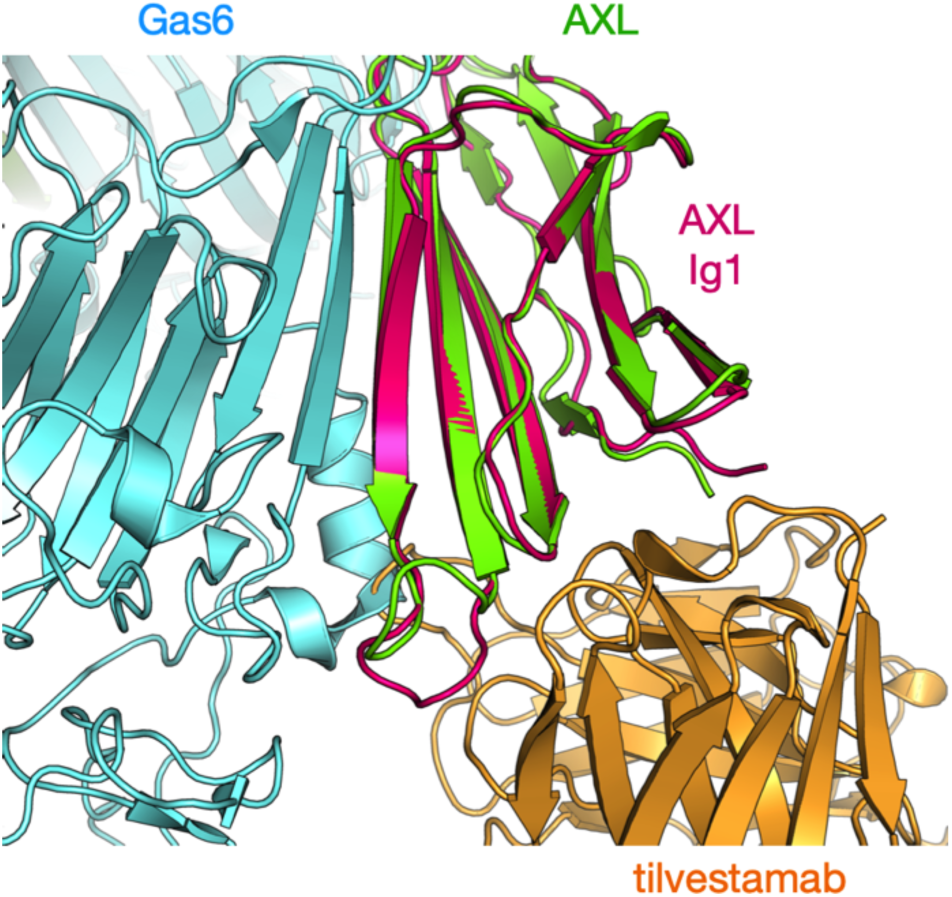
Superposition of the AlphaFold3 model (magenta/orange) and the AXL-GAS6 crystal structure (green/cyan). Note a small overlap in GAS6 and tilvestamab, which occupy binding surfaces close to each other on AXL Ig1.

## Conclusions

We have carried out an experimental screening of the CDR loops of tilvestamab to pinpoint key residues for AXL binding, and the AlphaFold3 model of the tilvestamab-AXL complex is in line with the mutagenesis data. Several interaction hotspots were identified, and one of the most interesting concerns the interactions of AXL residue Glu2, which is crucial for the interaction. The data point towards possibilities of affecting tilvestamab affinity further for various applications.

## Acknowledgements

We acknowledge the use of the Core Facility for Biophysics, Structural Biology, and Screening (BiSS) at the University of Bergen, which has received infrastructure funding from the Research Council of Norway (RCN) through NORCRYST (grant number 245828) and NOR–OPENSCREEN (grant number 245922). EC was supported by the Norwegian Research Council Industrial PhD Studentship 311399. We thank Hallvard Haugen for technical assistance.

## Abbreviations

BLI: biolayer interferometry
CDR: complementarity-determining region
FNIII: fibronectin type III
GAS6: growth arrest-specific protein 6
Ig: immunoglobulin
mAb: monoclonal antibody
osmY: osmotically inducible protein Y
scFv: single-chain variable fragment
TAM: TYRO3, AXL, MERTK
V_H_: heavy chain variable domain
V_L_: light chain variable domain

