## Supplemental data for "Paratope mapping of tilvestamab, an anti-AXL function blocking antibody, using high-throughput bacterial expression of secreted scFv-ompY fusion proteins"

**This PDF file includes:**

Supplemental Methods

Supplemental Figures 1-4

Supplemental Tables 1-2

#### **Supplemental Methods**

*Surface plasmon resonance analysis of binding of antibodies to AXL from different species*

Recombinant proteins in **Supplementary Figure 3A:** human AXL-Fc chimera (Evitria), recombinant mouse Axl-Fc chimera (R&D Systems 854-AX), and recombinant rhesus monkey AXL-Fc chimera were coupled to a CM5 sensor chip (Biacore BR-1000-14) using an amine coupling kit (Biacore BR-1000-50). Briefly, proteins were adjusted to 5 μg/ml in 10 mM sodium acetate pH 4.8 and the immobilisation protocol was run on a Biacore 3000 (GE Healthcare), aiming for 100 RU coupling. Chip performance was tested using antibody YW327.6S2var (Evitria) at 10 μg/ml in HBS-EP (Biacore 1001-88) with 1 min injection time and 20 μl/min flow rate for 10 cycles using 10 mM HCl and 1M NaCl as regeneration solution (30 s at 50μl/min followed by 1 min stabilisation). Binding of tilvestamab was tested similarly using 7.5 μg/ml antibody.

*Biolayer interferometry analysis of binding of antibodies to AXL variants*

Recombinant proteins in **Supplementary Figure 3C** were captured on streptavidin-coated dip and read biosensors (Forte-Bio 18-5019) by pre-treating the tips with 20 μg/ml biotin-conjugated donkey anti-human IgG Fc (Jackson ImmunoResearch 709-065-098) in PBS for 60 s. Tips were then saturated with antibody at 20 μg/ml in PBS for 300 s. A baseline was taken (60 s PBS) before a tip was immersed in each of the recombinant antigens and the association measured for 600 s. Tips were regenerated by three cycles of immersion in 10 mM glycine, pH 1.5 for 5 s followed by neutralisation in PBS for 5 s, before continuing with the next antibody. Antibodies were tested in the order tilvestamab then YW327.6S2var. For association with 1H12, the order of loading and association was reversed, as the mouse antibody 1H12 does not bind to the donkey anti-human IgG Fc. Recombinant proteins in **Supplementary Figure 4B** were dialysed to PBS and biotinylated with EZ-Link NHS-PEG4-Biotin (ThermoFisher 21329) by reconstituting the reagent at 20 mM in DMSO and diluting to 0.2 mM with water. 5 µl diluted biotinylation reagent was added to 1 ml recombinant protein, incubated for 30 min and dialysed extensively against PBS. Biotinylated recombinant proteins were captured on streptavidin-coated dip and read biosensors for 1200 s, a 600 s baseline was recorded before association with tilvestamab (10 µg/ml) or YW327.6S2var (6.6 µg/ml) for 100 s and dissociation in buffer for 600 s using fresh biosensors for each antibody.

Production of recombinant proteins for epitope mapping

Recombinant proteins were purchased from Evitria (Switzerland) or produced in-house by cloning the desired coding sequence into plasmid pCMV-3Tag-1A (Agilent). In each case, protein was produced by transient transfection of CHO cells followed by purification of the Fc-fusion proteins over a Protein A column (MAbSelect XTRA, GE Healthcare) with elution at low pH.

*Sequences of recombinant proteins and plasmids*

For proteins produced commercially, the final sequence is given, as the signal peptides used are proprietary. For proteins produced in-house, signal peptides are given in lower-case.

Human AXL Fc (AXL Fc, Evitria)

```
EESPFVGNPGNITGARGLTGTLRCQLQVQGEPPEVHWLRDGQILELADSTQTQVPLGEDEQDDWIVVSQLRITSL
QLSDTGQYQCLVFLGHQTFVSQPGYVGLEGLPYFLEEPEDRTVAANTPFNLSCQAQGPPEPVDLLWLQDAVPLAT
APGHGPQRSLHVPGLNKTSSFSCEAHNAKGVTTSRTATITVLPQQPRNLHLVSRQPTELEVAWTPGLSGIYPLTH
CTLQAVLSDDGMGIQAGEPDPPEEPLTSQASVPPHQLRLGSLHPHTPYHIRVACTSSQGPSSWTHWLPVETPEGV
PLGPPENISATRNGSQAFVHWQEPRAPLQGTLLGYRLAYQGQDTPEVLMDIGLRQEVTLELQGDGSVSNLTVCVA
AYTAAGDGPWSLPVPLEAWRPGQAQPVHQLVKEGGGGSGGGGSGGGGSDKTHTCPPCPAPELLGGPSVFLFPPKP
KDTLMISRTPEVTCVVVDVSHEDPEVKFNWYVDGVEVHNAKTKPREEQYNSTYRVVSVLTVLHQDWLNGKEYKCK
VSNKALPAPIEKTISKAKGQPREPQVYTLPPSREEMTKNQVSLTCLVKGFYPSDIAVEWESNGQPENNYKTTPPV
LDSDGSFFLYSKLTVDKSRWQQGNVFSCSVMHEALHNHYTQKSLSLSPGK
```

Rhesus Monkey AXL-Fc (RheAXL Fc, Evitria)

```
EESPFVGNPGNITGARGLTGTLRCQLQVQGEPPEVHWLRDGQILELADSTQTQVPLGEDEQDDWIVVSQLRIASL
QLSDAGQYQCLVFLGHQNFVSQPGYVGLEGLPYFLEEPEDRTVAANTHFNLSCQAQGPPEPVDLLWLQDAVPLAT
```

APGHGPQRNLHVPGLNKTSSFSCEAHNAKGVTTSRTATITVLPQQPRNLHLVSRQPTELEVAWTPGLSGIYPLTH
CTLQAMLSDNEVGIQAGEPDPPEEPLTLQASVPPHQLRLGSLHPHTPYHIRVACTSSQGPSSWTHWLPVETPEGV
PLGPPENISATRNGSQAFVHWQEPRAPLQGTLLGYRLAYQGQDTPEVLMDIGLRQEVTLELQGDGSVSNLTVCVA
AYTAAGDGPWSLPVPLEAWRPGQAQPVHQLVKEGGGGSGGGGSGGGGSDKTHTCPPCPAPELLGGPSVFLFPPKP
KDTLMISRTPEVTCVVVDVSHEDPEVKFNWYVDGVEVHNAKTKPREEQYNSTYRVVSVLTVLHQDWLNGKEYKCK
VSNKALPAPIEKTISKAKGQPREPQVYTLPPSREEMTKNQVSLTCLVKGFYPSDIAVEWESNGQPENNYKTTPPV
LDSDGSFFLYSKLTVDKSRWQQGNVFSCSVMHEALHNHYTQKSLSLSPGK

#### Mouse AXL-Fc (MmAXL Fc, RnD Systems)

HKDTQTEAGSPFVGNPGNITGARGLTGTLRCELQVQGEPPEVVWLRDGQILELADNTQTQVPLGEDWQDEWKVVS
QLRISALQLSDAGEYQCMVHLEGRTFVSQPGFVGLEGLPYFLEEPEDKAVPANTPFNLSCQAQGPPEPVTLLWLQ
DAVPLAPVTGHSSQHSLQTPGLNKTSSFSCEAHNAKGVTTSRTATITVLPQRPHHLHVVSRQPTELEVAWTPGLS
GIYPLTHCNLQAVLSDDGVGIWLGKSDPPEDPLTLQVSVPPHQLRLEKLLPHTPYHIRISCSSSQGPSPWTHWLP
VETTEGVPLGPPENVSAMRNGSQVLVRWQEPRVPLQGTLLGYRLAYRGQDTPEVLMDIGLTREVTLELRGDRPVA
NLTVSVTAYTSAGDGPWSLPVPLEPWRPGQGQPLHHLVSEPPPRAFSWPIEGRMDPEPRGPTIKPCPPCKCPAPN
LLGGPSVFIFPPKIKDVLMISLSPIVTCVVVDVSEDDPDVQISWFVNNVEVHTAQTQTHREDYNSTLRVVSALPI
QHQDWMSGKEFKCKVNNKDLPAPIERTISKPKGSVRAPQVYVLPPPEEEMTKKQVTLTCMVTDFMPEDIYVEWTN
NGKTELNYKNTEPVLDSDGSYFMYSKLRVEKKNWVERNSYSCSVVHEGLHNHHTTKSFSRTPGK

#### AXL IG1-Fc

mawrcprmgrvplawclalcgwacmaprgtqaEESPFVGNPGNITGARGLTGTLRCQLQVQGEPPEVHWLRDGQI
LELADSTQTQVPLGEDEQDDWIVVSQLRITSLQLSDTGQYQCLVFLGHQTFVSQPGYVGLEGSGGGGSGGGGSGG
GGSDKTHTCPPCPAPELLGGPSVFLFPPKPKDTLMISRTPEVTCVVVDVSHEDPEVKFNWYVDGVEVHNAKTKPR
EEQYNSTYRVVSVLTVLHQDWLNGKEYKCKVSNKALPAPIEKTISKAKGQPREPQVYTLPPSRDELTKNQVSLTC
LVKGFYPSDIAVEWESNGQPENNYKTTPPVLDSDGSFFLYSKLTVDKSRWQQGNVFSCSVMHEALHNHYTQKSLS
LSPGK

#### MsHsAxl-Fc

mgrvplawwlalccwgcaaHKDTQTEAGSPFVGNPGNITGARGLTGTLRCQLQVQGEPPEVHWLRDGQILELADS
TQTQVPLGEDEQDDWIVVSQLRITSLQLSDTGQYQCLVFLGHQTFVSQPGYVGLEGLPYFLEEPEDKAVPANTPF
NLSCQAQGPPEPVTLLWLQDAVPLAPVTGHSSQHSLQTPGLNKTSSFSCEAHNAKGVTTSRTATITVLPQRPHHL
HVVSRQPTELEVAWTPGLSGIYPLTHCNLQAVLSDDGVGIWLGKSDPPEDPLTLQVSVPPHQLRLEKLLPHTPYH
IRISCSSSQGPSPWTHWLPVETTEGVPLGPPENVSAMRNGSQVLVRWQEPRVPLQGTLLGYRLAYRGQDTPEVLM
DIGLTREVTLELRGDRPVANLTVSVTAYTSAGDGPWSLPVPLEPWRPGQGQPLHHLVSEPISGGGGSGGGGSGGG
GSDKTHTCPPCPAPELLGGPSVFLFPPKPKDTLMISRTPEVTCVVVDVSHEDPEVKFNWYVDGVEVHNAKTKPRE
EQYNSTYRVVSVLTVLHQDWLNGKEYKCKVSNKALPAPIEKTISKAKGQPREPQVYTLPPSRDELTKNQVSLTCL
VKGFYPSDIAVEWESNGQPENNYKTTPPVLDSDGSFFLYSKLTVDKSRWQQGNVFSCSVMHEALHNHYTQKSLSL
SPGK

#### MsHsAxl-Fc EAG>EE

mgrvplawwlalccwgcaaHKDTQTEESPFVGNPGNITGARGLTGTLRCQLQVQGEPPEVHWLRDGQILELADST
QTQVPLGEDEQDDWIVVSQLRITSLQLSDTGQYQCLVFLGHQTFVSQPGYVGLEGLPYFLEEPEDKAVPANTPFN
LSCQAQGPPEPVTLLWLQDAVPLAPVTGHSSQHSLQTPGLNKTSSFSCEAHNAKGVTTSRTATITVLPQRPHHLH
VVSRQPTELEVAWTPGLSGIYPLTHCNLQAVLSDDGVGIWLGKSDPPEDPLTLQVSVPPHQLRLEKLLPHTPYHI
RISCSSSQGPSPWTHWLPVETTEGVPLGPPENVSAMRNGSQVLVRWQEPRVPLQGTLLGYRLAYRGQDTPEVLMD
IGLTREVTLELRGDRPVANLTVSVTAYTSAGDGPWSLPVPLEPWRPGQGQPLHHLVSEPISGGGGSGGGGSGGGG
SDKTHTCPPCPAPELLGGPSVFLFPPKPKDTLMISRTPEVTCVVVDVSHEDPEVKFNWYVDGVEVHNAKTKPREE
QYNSTYRVVSVLTVLHQDWLNGKEYKCKVSNKALPAPIEKTISKAKGQPREPQVYTLPPSRDELTKNQVSLTCLV
KGFYPSDIAVEWESNGQPENNYKTTPPVLDSDGSFFLYSKLTVDKSRWQQGNVFSCSVMHEALHNHYTQKSLSLS
PGK

#### BGB289 pET-22b (+) (see **Supplemental Figure 1)**

```
FEATURES             Location/Qualifiers
     rep_origin      10..465
                     /note="f1 origin"
     CDS             597..1454
                     /note="bla AmpR"
     rep_origin      2215
                     /note="ColE1 pBR322 origin"
     CDS             complement(2646..2837)
                     /note="Rop"
     CDS             complement(3649..4728)
                     /note="lacI"
     regulatory      5115..5131
                     /regulatory_class="promoter"
                     /note=""
                     /note="T7 promoter"
     regulatory      5134..5158
                     /regulatory_class=""
                     /note="lac operator"
     CDS             5203..5286
                     /note="osmY signal peptide"
     misc_feature    5287..>5294
                     /note="SfiI/RI multiple cloning site"
     CDS             5299..6081
                     /note="Recombinant antibody fragment;"
     CDS             5302..5655
                     /note="H2L1 VH domain"
     CDS             5710..6045
                     /note="H2L1 VL domain"
     CDS             6082..6600
                     /locus_tag="SR36_22250"
                     /note="osmY"
                     /product="periplasmic protein"
     CDS             6601..6666
                     /note="Myc-His Tag"
     regulatory      6731..6777
                     /regulatory_class="terminator"
                     /note=""
                     /note="T7 terminator"
ORIGIN
        1 GCGAATGGGA CGCGCCCTGT AGCGGCGCAT TAAGCGCGGC GGGTGTGGTG GTTACGCGCA
       61 GCGTGACCGC TACACTTGCC AGCGCCCTAG CGCCCGCTCC TTTCGCTTTC TTCCCTTCCT
      121 TTCTCGCCAC GTTCGCCGGC TTTCCCCGTC AAGCTCTAAA TCGGGGGCTC CCTTTAGGGT
      181 TCCGATTTAG TGCTTTACGG CACCTCGACC CCAAAAAACT TGATTAGGGT GATGGTTCAC
      241 GTAGTGGGCC ATCGCCCTGA TAGACGGTTT TTCGCCCTTT GACGTTGGAG TCCACGTTCT
      301 TTAATAGTGG ACTCTTGTTC CAAACTGGAA CAACACTCAA CCCTATCTCG GTCTATTCTT
      361 TTGATTTATA AGGGATTTTG CCGATTTCGG CCTATTGGTT AAAAAATGAG CTGATTTAAC
      421 AAAAATTTAA CGCGAATTTT AACAAAATAT TAACGTTTAC AATTTCAGGT GGCACTTTTC
      481 GGGGAAATGT GCGCGGAACC CCTATTTGTT TATTTTTCTA AATACATTCA AATATGTATC
      541 CGCTCATGAG ACAATAACCC TGATAAATGC TTCAATAATA TTGAAAAAGG AAGAGTATGA
      601 GTATTCAACA TTTCCGTGTC GCCCTTATTC CCTTTTTTGC GGCATTTTGC CTTCCTGTTT
      661 TTGCTCACCC AGAAACGCTG GTGAAAGTAA AAGATGCTGA AGATCAGTTG GGTGCACGAG
      721 TGGGTTACAT CGAACTGGAT CTCAACAGCG GTAAGATCCT TGAGAGTTTT CGCCCCGAAG
      781 AACGTTTTCC AATGATGAGC ACTTTTAAAG TTCTGCTATG TGGCGCGGTA TTATCCCGTA
      841 TTGACGCCGG GCAAGAGCAA CTCGGTCGCC GCATACACTA TTCTCAGAAT GACTTGGTTG
      901 AGTACTCACC AGTCACAGAA AAGCATCTTA CGGATGGCAT GACAGTAAGA GAATTATGCA
      961 GTGCTGCCAT AACCATGAGT GATAACACTG CGGCCAACTT ACTTCTGACA ACGATCGGAG
     1021 GACCGAAGGA GCTAACCGCT TTTTTGCACA ACATGGGGGA TCATGTAACT CGCCTTGATC
     1081 GTTGGGAACC GGAGCTGAAT GAAGCCATAC CAAACGACGA GCGTGACACC ACGATGCCTG
     1141 CAGCAATGGC AACAACGTTG CGCAAACTAT TAACTGGCGA ACTACTTACT CTAGCTTCCC
     1201 GGCAACAATT AATAGACTGG ATGGAGGCGG ATAAAGTTGC AGGACCACTT CTGCGCTCGG
     1261 CCCTTCCGGC TGGCTGGTTT ATTGCTGATA AATCTGGAGC CGGTGAGCGT GGGTCTCGCG
     1321 GTATCATTGC AGCACTGGGG CCAGATGGTA AGCCCTCCCG TATCGTAGTT ATCTACACGA
     1381 CGGGGAGTCA GGCAACTATG GATGAACGAA ATAGACAGAT CGCTGAGATA GGTGCCTCAC
     1441 TGATTAAGCA TTGGTAACTG TCAGACCAAG TTTACTCATA TATACTTTAG ATTGATTTAA
     1501 AACTTCATTT TTAATTTAAA AGGATCTAGG TGAAGATCCT TTTGATAAT CTCATGACCA
```

1561 AAATCCCTTA ACGTGAGTTT TCGTTCCACT GAGCGTCAGA CCCCGTAGAA AAGATCAAAG
1621 GATCTTCTTG AGATCCTTTT TTTCTGCGCG TAATCTGCTG CTTGCAAACA AAAAAACCAC
1681 CGCTACCAGC GGTGGTTTGT TTGCCGGATC AAGAGCTACC AACTCTTTTT CCGAAGGTAA
1741 CTGGCTTCAG CAGAGCGCAG ATACCAAATA CTGTCCTTCT AGTGTAGCCG TAGTTAGGCC
1801 ACCACTTCAA GAACTCTGTA GCACCGCCTA CATACCTCGC TCTGCTAATC CTGTTACCAG
1861 TGGCTGCTGC CAGTGGCGAT AAGTCGTGTC TTACCGGGTT GGACTCAAGA CGATAGTTAC
1921 CGGATAAGGC GCAGCGGTCG GGCTGAACGG GGGGTTCGTG CACACAGCCC AGCTTGGAGC
1981 GAACGACCTA CACCGAACTG AGATACCTAC AGCGTGAGCT ATGAGAAAGC GCCACGCTTC
2041 CCGAAGGGAG AAAGGCGGAC AGGTATCCGG TAAGCGGCAG GGTCGGAACA GGAGAGCGCA
2101 CGAGGGAGCT TCCAGGGGGA AACGCCTGGT ATCTTTATAG TCCTGTCGGG TTTCGCCACC
2161 TCTGACTTGA GCGTCGATTT TTGTGATGCT CGTCAGGGGG GCGGAGCCTA TGGAAAAACG
2221 CCAGCAACGC GGCCTTTTTA CGGTTCCTGG CCTTTTGCTG GCCTTTTGCT CACATGTTCT
2281 TTCCTGCGTT ATCCCCTGAT TCTGTGGATA ACCGTATTAC CGCCTTTGAG TGAGCTGATA
2341 CCGCTCGCCG CAGCCGAACG ACCGAGCGCA GCGAGTCAGT GAGCGAGGAA GCGGAAGAGC
2401 GCCTGATGCG GTATTTTCTC CTTACGCATC TGTGCGGTAT TTCACACCGC ATATATGGTG
2461 CACTCTCAGT ACAATCTGCT CTGATGCCGC ATAGTTAAGC CAGTATACAC TCCGCTATCG
2521 CTACGTGACT GGGTCATGGC TGCGCCCCGA CACCCGCCAA CACCCGCTGA CGCGCCCTGA
2581 CGGGCTTGTC TGCTCCCGGC ATCCGCTTAC AGACAAGCTG TGACCGTCTC CGGGAGCTGC
2641 ATGTGTCAGA GGTTTTCACC GTCATCACCG AAACGCGCGA GGCAGCTGCG GTAAAGCTCA
2701 TCAGCGTGGT CGTGAAGCGA TTCACAGATG TCTGCCTGTT CATCCGCGTC CAGCTCGTTG
2761 AGTTTCTCCA GAAGCGTTAA TGTCTGGCTT CTGATAAAGC GGGCCATGTT AAGGGCGGTT
2821 TTTTCCTGTT TGGTCACTGA TGCCTCCGTG TAAGGGGGAT TTCTGTTCAT GGGGGTAATG
2881 ATACCGATGA AACGAGAGAG GATGCTCACG ATACGGGTTA CTGATGATGA ACATGCCCGG
2941 TTACTGGAAC GTTGTGAGGG TAAACAACTG GCGGTATGGA TGCGGCGGGA CCAGAGAAAA
3001 ATCACTCAGG GTCAATGCCA GCGCTTCGTT AATACAGATG TAGGTGTTCC ACAGGGTAGC
3061 CAGCAGCATC CTGCGATGCA GATCCGGAAC ATAATGGTGC AGGGCGCTGA CTTCCGCGTT
3121 TCCAGACTTT ACGAAACACG GAAACCGAAG ACCATTCATG TTGTTGCTCA GGTCGCAGAC
3181 GTTTTGCAGC AGCAGTCGCT TCACGTTCGC TCGCGTATCG GTGATTCATT CTGCTAACCA
3241 GTAAGGCAAC CCCGCCAGCC TAGCCGGGTC CTCAACGACA GGAGCACGAT CATGCGCACC
3301 CGTGGGGCCG CCATGCCGGC GATAATGGCC TGCTTCTCGC CGAAACGTTT GGTGGCGGGA
3361 CCAGTGACGA AGGCTTGAGC GAGGGCGTGC AAGATTCCGA ATACCGCAAG CGACAGGCCG
3421 ATCATCGTCG CGCTCCAGCG AAAGCGGTCC TCGCCGAAAA TGACCCAGAG CGCTGCCGGC
3481 ACCTGTCCTA CGAGTTGCAT GATAAAGAAG ACAGTCATAA GTGCGGCGAC GATAGTCATG
3541 CCCCGCGCCC ACCGGAAGGA GCTGACTGGG TTGAAGGCTC TCAAGGGCAT CGGTCGAGAT
3601 CCCGGTGCCT AATGAGTGAG CTAACTTACA TTAATTGCGT TGCGCTCACT GCCCGCTTTC
3661 CAGTCGGGAA ACCTGTCGTG CCAGCTGCAT TAATGAATCG GCCAACGCGC GGGGAGAGGC
3721 GGTTTGCGTA TTGGGCGCCA GGGTGGTTTT TCTTTTCACC AGTGAGACGG GCAACAGCTG
3781 ATTGCCCTTC ACCGCCTGGC CCTGAGAGAG TTGCAGCAAG CGGTCCACGC TGGTTTGCCC
3841 CAGCAGGCGA AAATCCTGTT TGATGGTGGT TAACGGCGGG ATATAACATG AGCTGTCTTC
3901 GGTATCGTCG TATCCCACTA CCGAGATATC CGCACCAACG CGCAGCCCGG ACTCGGTAAT
3961 GGCGCGCATT GCGCCCAGCG CCATCTGATC GTTGGCAACC AGCATCGCAG TGGGAACGAT
4021 GCCCTCATTC AGCATTTGCA TGGTTTGTTG AAAACCGGAC ATGGCACTCC AGTCGCCTTC
4081 CCGTTCCGCT ATCGGCTGAA TTTGATTGCG AGTGAGATAT TTATGCCAGC CAGCCAGACG
4141 CAGACGCGCC GAGACAGAAC TTAATGGGCC CGCTAACAGC GCGATTTGCT GGTGACCCAA
4201 TGCGACCAGA TGCTCCACGC CCAGTCGCGT ACCGTCTTCA TGGGAGAAAA TAATACTGTT
4261 GATGGGTGTC TGGTCAGAGA CATCAAGAAA TAACGCCGGA ACATTAGTGC AGGCAGCTTC
4321 CACAGCAATG GCATCCTGGT CATCCAGCGG ATAGTTAATG ATCAGCCCAC TGACGCGTTG
4381 CGCGAGAAGA TTGTGCACCG CCGCTTTACA GGCTTCGACG CCGCTTCGTT CTACCATCGA
4441 CACCACCACG CTGGCACCCA GTTGATCGGC GCGAGATTTA ATCGCCGCGA CAATTTGCGA
4501 CGGCGCGTGC AGGGCCAGAC TGGAGGTGGC AACGCCAATC AGCAACGACT GTTTGCCCGC
4561 CAGTTGTTGT GCCACGCGGT TGGGAATGTA ATTCAGCTCC GCCATCGCCG CTTCCACTTT
4621 TTCCCGCGTT TTCGCAGAAA CGTGGCTGGC CTGGTTCACC ACGCGGGAAA CGGTCTGATA
4681 AGAGACACCG GCATACTCTG CGACATCGTA TAACGTTACT GGTTTCACAT TCACCACCCT
4741 GAATTGACTC TCTTCCGGGC GCTATCATGC CATACCGCGA AAGGTTTTGC GCCATTCGAT
4801 GGTGTCCGGG ATCTCGACGC TCTCCCTTAT GCGACTCCTG CATTAGGAAG CAGCCCAGTA
4861 GTAGGTTGAG GCCGTTGAGC ACCGCCGCCG CAAGGAATGG TGCATGCAAG GAGATGGCGC
4921 CCAACAGTCC CCCGGCCACG GGGCCTGCCA CCATACCCAC GCCGAAACAA GCGCTCATGA
4981 GCCCGAAGTG GCGAGCCCGA TCTTCCCCAT CGGTGATGTC GGCGATATAG GCGCCAGCAA
5041 CCGCACCTGT GGCGCCGGTG ATGCCGGCCA CGATGCGTCC GGCGTAGAGG ATCGAGATCT
5101 CGATCCCGCG AAATTAATAC GACTCACTAT AGGGGAATTG TGAGCGGATA ACAATTCCCC
5161 TCTAGAAATA ATTTTGTTTA ACTTTAAGAA GGAGATATAC ATATGACcAT GACccGtGTG
5221 AAGATTagcA AgACcCTGCT GGCgGTtATG cTGACCagcG CgGTgGCGAC CGGtagcGCg
5281 TAtGCGGGCC CAGCCGGCCT GGAAGTGCAG CTGGTTGAAA GCGGTGGCGG TCTGGTGCAA
5341 CCGGGCGGTA GCCTGCGTCT GAGCTGCGCG GCGAGCGGTT ACAGCTTCAC CGACTTTTAT
5401 ATCAACTGGG TGCGTCAGGC GCCGGGTAAA GGTCTGGAGT GGGTTGCGCG TATTTTCCCG
5461 GGCGGTGACA ACACCTACTA TAACGAAAAG TTCAAAGGTC GTTTTACCCT GAGCGCGGAT

5521 ACCAGCAAGA GCACCGCGTA CCTGCAGATG AACAGCCTGC GTGCGGAGGA CACCGCGGTT
5581 TACTATTGCG CGCGTCGTGG CCTGTACTAT GCGATGGATT ATTGGGGCCA AGGTACCCTG
5641 GTGACCGTTA GCAGCGCGAA AACCACCCCT CCTAAGCTTG AGGAAGGTGA ATTCAGCGAG
5701 GCACGCGTAG ACATCCAGAT GACCCAGAGC CCGAGCAGCC TGAGCGCGAG CGTGGGCGAT
5761 CGTGTTACCA TCACCTGCCG TAGCAGCCAG AGCCTGGTTC ACAGCAACGG TATTCCGTAC
5821 CTGCACTGGT ATCAGCAAAA GCCGGGCAAA GCGCCGAAGC TGCTGATCTA CCGTGTGAGC
5881 AACCGTTTCA GCGGTGTTCC GAGCCGTTTT AGCGGTAGCG GTAGCGGTAC CGACTTCACC
5941 CTGACCATTA GCAGCCTGCA ACCGGAGGAT TTTGCGACCT ACTATTGCAG CCAGGGTACC
6001 CATGTGCCGC CGACCTTCGG TCAAGGCACC AAAGTTGAAA TCAAGCGTGC GGATGCGGCG
6061 CCGACCGTGT CTGCGGCCGC TGAGAACAAC GCGCAGACCA CCAACGAAAG CGCGGGTCAA
6121 AAAGTGGATA GCAGCATGAA CAAGGTTGGC AACTTCATGG ACGATAGCGC GATTACCGCG
6181 AAGGTGAAAG CGGCGCTGGT TGACCACGAT AACATCAAAA GCACCGACAT TAGCGTGAAA
6241 ACCGATCAGA AGGTGGTTAC CCTGAGCGGT TTTGTTGAGA GCCAGGCGCA AGCGGAGGAA
6301 GCGGTGAAAG TTGCGAAGGG TGTGGAAGGC GTTACCAGCG TGAGCGACAA ACTGCACGTG
6361 CGTGATGCGA AAGAGGGTAG CGTTAAAGGT TATGCGGGTG ACACCGCGAC CACCAGCGAA
6421 ATCAAGGCGA AACTGCTGGC GGACGATATT GTGCCGAGCC GTCACGTGAA GGTTGAAACC
6481 ACCGACGGTG TGGTTCAACT GAGCGGCACC GTTGACAGCC AGGCGCAAAG CGATCGTGCG
6541 GAAAGCATTG CGAAAGCGGT TGATGGTGTG AAGAGCGTTA AAAACGACCT GAAAACCAAG
6601 GGATCCGAAC AGAAACTGAT TAGCGAAGAG GACCTGAGCC TCGAGCACCA CCACCACCAC
6661 CACTGAGATC CGGCTGCTAA CAAAGCCCGA AAGGAAGCTG AGTTGGCTGC TGCCACCGCT
6721 GAGCAATAAC TAGCATAACC CCTTGGGGCC TCTAAACGGG TCTTGAGGGG TTTTTTGCTG
6781 AAAGGAGGAA CTATATCCGG ATtg

#### Supplemental Figures

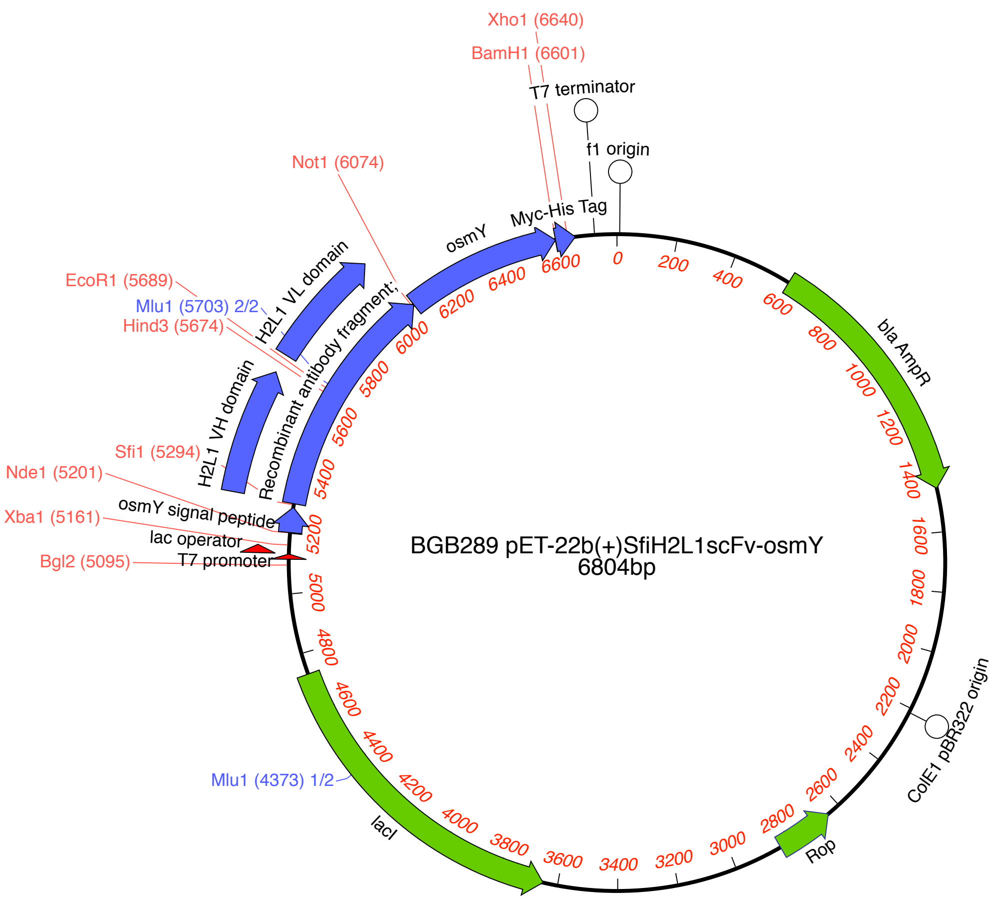


**Supplemental Figure 1. Parental scFv-ompY expression vector.**

```
        1          11         21         31         41         51
AXL Ig1 EESPFVGNPG NITGARGLTG TLRCQLQVQG EPPEVHWLRD GQILELADST QTQVPLGEDE
        61         71         81         91
        QDDWIVVSQL RITSLQLSDT GQYQCLVFLG HQTFVS
```

**Supplemental Figure 2. Residue numbering for the mature AXL Ig1.** Amino acid positions are indicated with numbers above the sequence. Due to omission of the N-terminus signal peptide from the mature protein sequence, the residue numbering starts from position E33 of the original numbering scheme by Bryan et al.,[1] which is considered E1 in the context of this study.

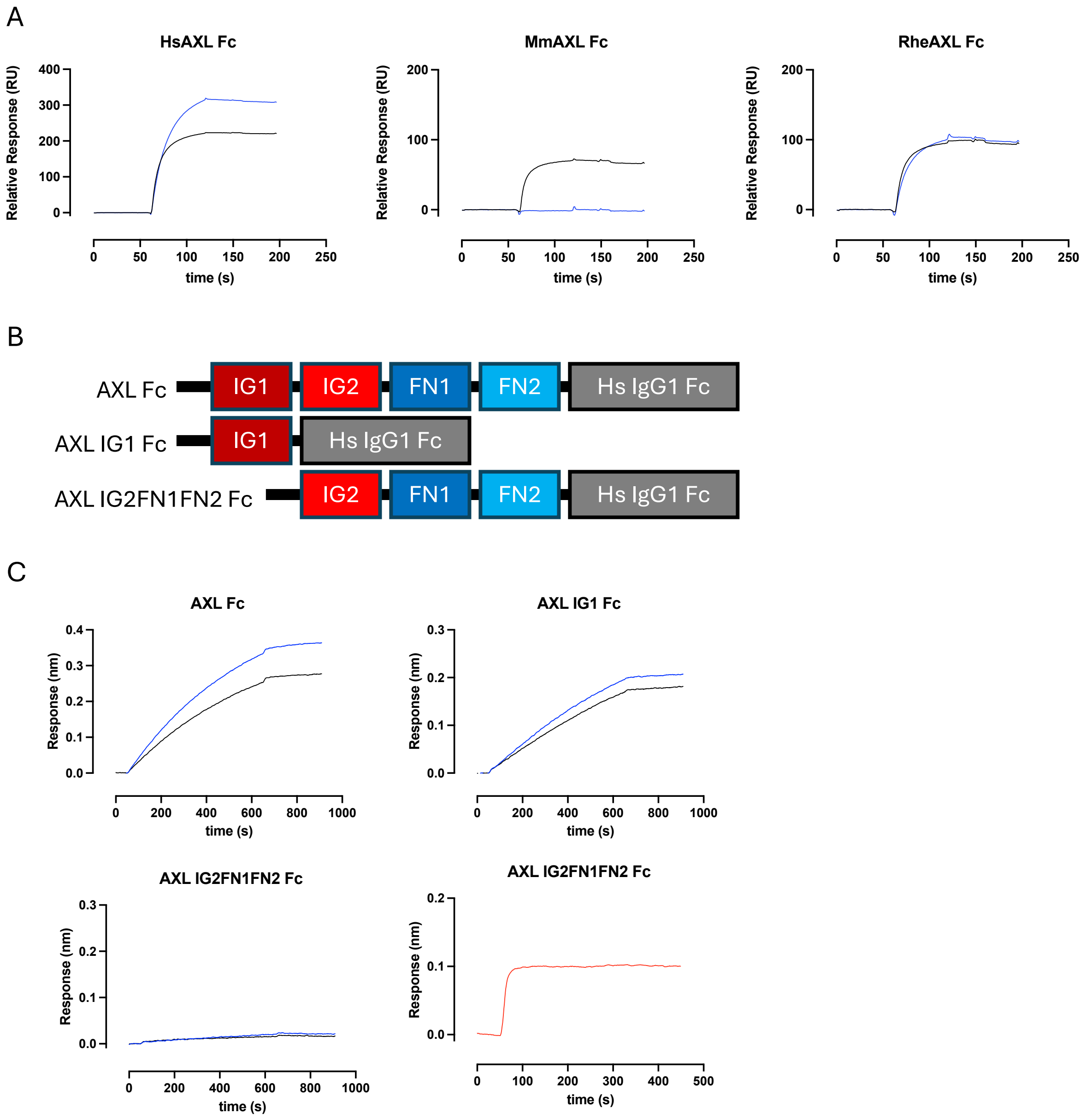


**Supplementary Figure 3. Tilvestamab binds to the Ig1 domain of human AXL. (A)** Surface plasmon resonance sensorgrams showing tilvestamab binding (blue) to human AXL Fc (left panel) and rhesus monkey AXL Fc (right panel) but not to mouse AXL Fc (middle panel) immobilised on a Biacore CM5 sensor chip. Binding of an anti-AXL monoclonal antibody that recognises human, rhesus monkey, and mouse AXL (YW327.6S2var)[2] is also shown as a control for the presence and integrity of the immobilised proteins (black). **(B)** Structural composition of the recombinant proteins produced to map tilvestamab epitope. **(C)** BLI sensograms of association of tilvestamab (blue) to immobilised recombinant proteins. Tilvestamab binds recombinant proteins incorporating the entire AXL extracellular domain (left upper panel) or just the Ig1 domain (right upper panel) but does not bind a protein lacking the Ig1 domain (lower panels). YW327.6S2var (black) and 1H12, an anti-AXL monoclonal antibody that binds the Ig2 domain of human AXL (red)[3] were used as controls for the presence and integrity of recombinant proteins not bound by tilvestamab.

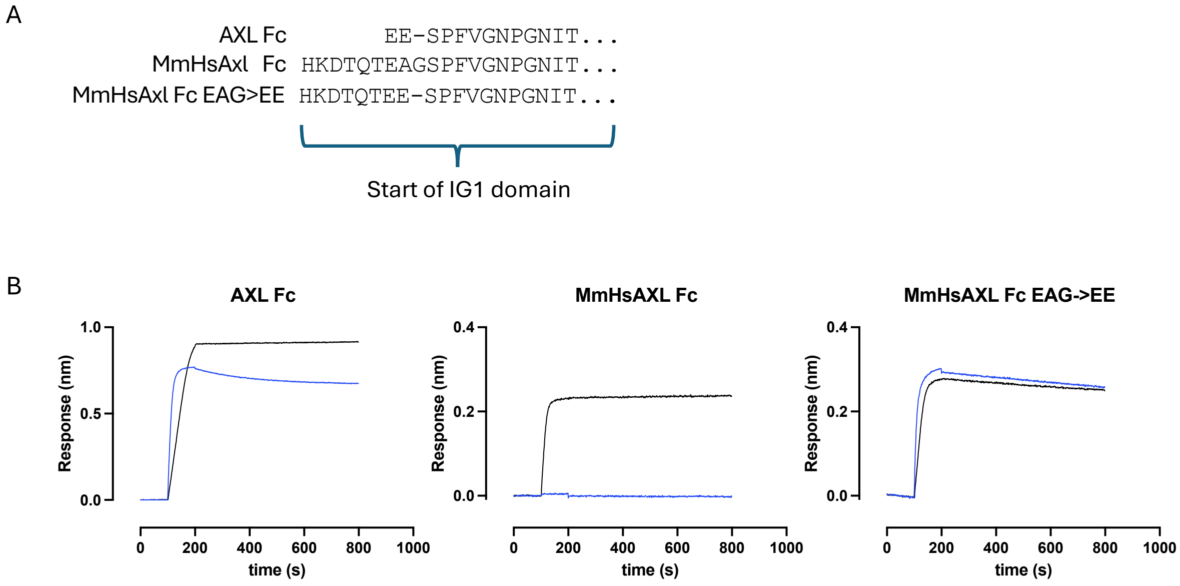


**Supplementary Figure 3. A single glutamate residue is essential for tilvestamab binding to AXL. (A)** Comparison of the N-terminal sequences of human AXL Fc, and mouse Axl Fc with a humanized Ig1 domain with and without mutation of the mouse N-terminal peptide from HKDTQTEAG (MmHsAxl) to HKDTQTEE (MmHsAxl Fc EAG>EE). Sequences are shown from the first amino-acid following the predicted signal peptide cleave sites for human and mouse AXL. In each case the sequence of the Ig1 domain is identical. **(B)** BLI sensograms of association of tilvestamab (blue) or YW327.6S2var (black) to immobilised recombinant proteins. Tilvestamab is unable to bind the Ig1 domain in the context of the mouse N-terminal peptide (middle panel). Mutation of the mouse sequence EAG to match the human sequence EE restores binding (right panel).

### Supplemental Tables

#### Supplemental Table 1. Primers for generation of mutations in heavy chain CDRs

| Name | Sequence |
|---|---|
| **VH CDR1** | |
| VHCDR1_new f | TGGGTGCGTCAGGCGCCGGGTA |
| VHCDR1_Parentr | GTT GAT ATA AAA GTC GGT GAA GCT GTA ACC GCT CG<br><N I Y F D T F..... |
| VHCDR1_N36Ar | Ggc GAT ATA AAA GTC GGT GAA GCT GTA ACC GCT CG<br><**A** I Y F D T F..... |
| VHCDR1_I35Ar | GTT Ggc ATA AAA GTC GGT GAA GCT GTA ACC GCT CG<br><N **A** Y F D T F..... |
| VHCDR1_Y34Ar | GTT GAT Agc AAA GTC GGT GAA GCT GTA ACC GCT CG<br><N I **A** F D T F..... |
| VHCDR1_F33Ar | GTT GAT ATA Agc GTC GGT GAA GCT GTA ACC GCT CG<br><N I Y **A** D T F..... |
| VHCDR1_Y34Xr | GTT GAT Akn AAA GTC GGT GAA GCT GTA ACC GCT CG<br><N I **X** F D T F..... |
| VHCDR1_F33Xr | GTT GAT ATA Akn GTC GGT GAA GCT GTA ACC GCT CG<br><N I Y **X** D T F..... |
| VHCDR1_D32Ar | GTT GAT ATA AAA GgC GGT GAA GCT GTA ACC GCT CG<br><N I Y F **A** T F..... |
| VHCDR1_T31Ar | GTT GAT ATA AAA GTC GGc GAA GCT GTA ACC GCT CG<br><N I Y F D **A** F..... |
| **Additional oligos for gap-filling:** | |
| VHCDR1_F33Dr | GTT GAT ATA Atc GTC GGT GAA GCT GTA ACC GCT CG |
| VHCDR1_F33Tr | GTT GAT ATA Agt GTC GGT GAA GCT GTA ACC GCT CG |
| VHCDR1_Y34Dr | GTT GAT Atc AAA GTC GGT GAA GCT GTA ACC GCT CG |
| VHCDR1_Y34Sr | GTT GAT Act AAA GTC GGT GAA GCT GTA ACC GCT CG |
| **VH CDR2 RHS** | |
| VHCDR2RHS_new r | GTGTTGTCACCGCCCGGGAA |
| VHCDR2RHS_Parent | CTA CTA TAA CGA AAA GTT CAA AGG TCG TTT TACC<br>Y Y N E K F K G R F T |
| VHCDR2RHS_Y60Af n | C GCC TAT AAC GAA AAG TTC AAA GGT CGT TTT ACC<br>**A** Y N E K F K G R F T |
| VHCDR2RHS_Y60Xf n | C NMC TAT AAC GAA AAG TTC AAA GGT CGT TTT ACC<br>**X** Y N E K F K G R F T |
| VHCDR2RHS_Y61Af n | C TAC GCT AAC GAA AAG TTC AAA GGT CGT TTT ACC<br>Y **A** N E K F K G R F T |
| VHCDR2RHS_N62Af n | C TAC TAT GCC GAA AAG TTC AAA GGT CGT TTT ACC<br>Y Y **A** E K F K G R F T |
| VHCDR2RHS_E63Af n | C TAC TAT AAC GCA AAG TTC AAA GGT CGT TTT ACC<br>Y Y N **A** K F K G R F T |
| VHCDR2RHS_K64Af n | C TAC TAT AAC GAA GCG TTC AAA GGT CGT TTT ACC<br>Y Y N E **A** F K G R F T |
| VHCDR2RHS_F65Af n | C TAC TAT AAC GAA AAG GCC AAA GGT CGT TTT ACC C<br>Y Y N E K **A** K G R F T |
| VHCDR2RHS_K66Af n | C TAC TAT AAC GAA AAG TTC GCA GGT CGT TTT ACC<br>Y Y N E K F **A** G R F T |
| VHCDR2RHS_G67Af n | C TAC TAT AAC GAA AAG TTC AAA GCT CGT TTT ACC<br>Y Y N E K F K **A** R F T |
| **Additional oligos for gap-filling:** | |
| VHCDR2RHS_Y60Pf | C cTC TAT AAC GAA AAG TTC AAA GGT CGT TTT ACC |
| **VH CDR2 LHS** | |
| VHCDR2LHS_r: | CGCAACCCACTCCAGACC, Tm= 68°C, Ta = 70°C |
| VHCDR2LHS_Parent | CGT ATT TTC CCG GGC GGT GAC AAC ACC TAC TAT AAC GA<br>R I F P G G D N T Y Y N E |
| VHCDR2LHS_R51Af | GCT ATT TTC CCG GGC GGT GAC AAC ACC TAC TAT AAC GA<br>**A** I F P G G D N T Y Y N E |
| VHCDR2LHS_I52Af | CGT GCT TTC CCG GGC GGT GAC AAC ACC TAC TAT AAC GA<br>R **A** F P G G D N T Y Y N E |
| VHCDR2LHS_F53Af | CGT ATT GCC CCG GGC GGT GAC AAC ACC TAC TAT AAC GA<br>R I **A** P G G D N T Y Y N E |
| VHCDR2LHS_P54Af | CGT ATT TTC GCG GGC GGT GAC AAC ACC TAC TAT AAC GA<br>R I F **A** G G D N T Y Y N E |
| VHCDR2LHS_G55Af | CGT ATT TTC CCG GCC GGT GAC AAC ACC TAC TAT AAC GA<br>R I F P **A** G D N T Y Y N E |
| VHCDR2LHS_G56Af | CGT ATT TTC CCG GGC GCT GAC AAC ACC TAC TAT AAC GA<br>R I F P G **A** D N T Y Y N E |
| VHCDR2LHS_D57Af | CGT ATT TTC CCG GGC GGT GCC AAC ACC TAC TAT AAC GA<br>R I F P G G **A** N T Y Y N E |
| VHCDR2LHS_N58Af | CGT ATT TTC CCG GGC GGT GAC GCC ACC TAC TAT AAC GA<br>R I F P G G D **A** T Y Y N E |
| VHCDR2LHS_T58Af | CGT ATT TTC CCG GGC GGT GAC AAC GCC TAC TAT AAC GA<br>R I F P G G D N **A** Y Y N E |
| VHCDR2LHS_R51Xf | NMT ATT TTC CCG GGC GGT GAC AAC ACC TAC TAT AAC GA<br>**X** I F P G G D N T Y Y N E |
| VHCDR2LHS_F53Xf | CGT ATT NMT CCG GGC GGT GAC AAC ACC TAC TAT AAC GA<br>R I **X** P G G D N T Y Y N E |
| **Additional oligos for gap-filling:** | |
| VHCDR2LHS_F53HYf | CGT ATT yaT CCG GGC GGT GAC AAC ACC TAC TAT AAC GA |

| Name | Sequence |
|---|---|
| **VH CDR3** | |
| VHCDR3_r | ACGCGCGCAATAGTAAAC, Tm=64°C, Ta=65 |
| VHCDR3_Parentf | CGT GGC CTG TAC TAT GCG ATG GAT TAT TGG GGC CAA G<br>R G L Y Y A M D Y W G Q |
| VHCDR3_R100A | gcT GGC CTG TAC TAT GCG ATG GAT TAT TGG GGC CAA G<br>A G L Y Y A M D Y W G Q |
| VHCDR3_G101A | CGT GcC CTG TAC TAT GCG ATG GAT TAT TGG GGC CAA G<br>R A L Y Y A M D Y W G Q |
| VHCDR3_L102A | CGT GGC gcG TAC TAT GCG ATG GAT TAT TGG GGC CAA G<br>R G A Y Y A M D Y W G Q |
| VHCDR3_Y103A | CGT GGC CTG gcC TAT GCG ATG GAT TAT TGG GGC CAA G<br>R G L A Y A M D Y W G Q |
| VHCDR3_Y104A | CGT GGC CTG TAC gcT GCG ATG GAT TAT TGG GGC CAA G<br>R G L Y a A M D Y W G Q |
| VHCDR3_M106A | CGT GGC CTG TAC TAT GCG gcg GAT TAT TGG GGC CAA G<br>R G L Y Y A A D Y W G Q |
| VHCDR3_D107A | CGT GGC CTG TAC TAT GCG ATG gcT TAT TGG GGC CAA G<br>R G L Y Y A M A Y W G Q |
| VHCDR3_Y108A | CGT GGC CTG TAC TAT GCG ATG GAT gcT TGG GGC CAA G<br>R G L Y Y A M D A W G Q |
| VHCDR3_R100X | NMT GGC CTG TAC TAT GCG ATG GAT TAT TGG GGC CAA G<br>X G L Y Y A M D Y W G Q |
| VHCDR3_L102X | CGT GGC NMT TAC TAT GCG ATG GAT TAT TGG GGC CAA G<br>R G X Y Y A M D Y W G Q |
| VHCDR3_Y103X | CGT GGC CTG NMT TAT GCG ATG GAT TAT TGG GGC CAA G<br>R G L X Y A M D Y W G Q |
| VHCDR3_Y104X | CGT GGC CTG TAC NMT GCG ATG GAT TAT TGG GGC CAA G<br>R G L Y X A M D Y W G Q |
| **Additional oligos for gap-filling:** | |
| VHCDR3_R100Y | taT GGC CTG TAC TAT GCG ATG GAT TAT TGG GGC CAA G |
| VHCDR3_Y103N | CGT GGC CTG aaT TAT GCG ATG GAT TAT TGG GGC CAA G |
| VHCDR3_Y103ST | CGT GGC CTG wcT TAT GCG ATG GAT TAT TGG GGC CAA G |
| VHCDR3_Y104P | CGT GGC CTG TAC ccT GCG ATG GAT TAT TGG GGC CAA G |

### Supplemental Table 2. Primers for generation of mutations in light chain CDRs

| Name | Sequence |
|---|---|
| **VL CDR1 LHS** | |
| VLCDR1LHS_r | GCAGGTGATGGTAACACGATC, Tm=66°C, Ta=67 |
| VLCDR1LHS_Parent f | CGT AGC AGC CAG AGC CTG GTT CAC AGC AAC GGT A<br>R S S Q S L V H S N G |
| VLCDR1LHS_R161A | gcT AGC AGC CAG AGC CTG GTT CAC AGC AAC GGT A<br>**A** S S Q S L V H S N G |
| VLCDR1LHS_S162A | CGT gcC AGC CAG AGC CTG GTT CAC AGC AAC GGT A<br>R **A** S Q S L V H S N G |
| VLCDR1LHS_S163A | CGT AGC gcC CAG AGC CTG GTT CAC AGC AAC GGT A<br>R S **A** Q S L V H S N G |
| VLCDR1LHS_Q164A | CGT AGC AGC GcG AGC CTG GTT CAC AGC AAC GGT A<br>R S S **A** S L V H S N G |
| VLCDR1LHS_S165A | CGT AGC AGC CAG gcC CTG GTT CAC AGC AAC GGT A<br>R S S Q **A** L V H S N G |
| VLCDR1LHS_L166A | CGT AGC AGC CAG AGC gcG GTT CAC AGC AAC GGT A<br>R S S Q S **A** V H S N G |
| VLCDR1LHS_V167A | CGT AGC AGC CAG AGC CTG GcT CAC AGC AAC GGT A<br>R S S Q S L **A** H S N G |
| VLCDR1LHS_H168A | CGT AGC AGC CAG AGC CTG GTT gcC AGC AAC GGT A<br>R S S Q S L V **A** S N G |
| VLCDR1LHS_H168X | CGT AGC AGC CAG AGC CTG GTT NMT AGC AAC GGT A<br>R S S Q S L V **X** S N G |
| Additional oligos for gap-filling: | |
| VLCDR1LHS_H168ST | CGT AGC AGC CAG AGC CTG GTT wcT AGC AAC GGT A |

| Name | Sequence |
|---|---|
| **VL CDR1 RHS** | |
| VLCDR1RHS_r | GTGAACCAGGCTCTGGCT, Tm=67°C, Ta=64 |
| VLCDR1RHS_Parent f | AGC AAC GGT ATT CCG TAC CTG CAC TGG TAT CAG C<br>S N G I P Y L H W Y Q |
| VLCDR1RHS_S169A | gcC AAC GGT ATT CCG TAC CTG CAC TGG TAT CAG C<br>**A** N G I P Y L H W Y Q |
| VLCDR1RHS_N170A | AGC gcC GGT ATT CCG TAC CTG CAC TGG TAT CAG C<br>S **A** G I P Y L H W Y Q |
| VLCDR1RHS_G171A | AGC AAC GcT ATT CCG TAC CTG CAC TGG TAT CAG C<br>S N **A** I P Y L H W Y Q |
| VLCDR1RHS_I172A | AGC AAC GGT gcT CCG TAC CTG CAC TGG TAT CAG C<br>S N G **A** P Y L H W Y Q |
| VLCDR1RHS_P173A | AGC AAC GGT ATT gcG TAC CTG CAC TGG TAT CAG C<br>S N G I **A** Y L H W Y Q |
| VLCDR1RHS_Y174A | AGC AAC GGT ATT CCG gcC CTG CAC TGG TAT CAG C<br>S N G I P **A** L H W Y Q |
| VLCDR1RHS_L175A | AGC AAC GGT ATT CCG TAC gcG CAC TGG TAT CAG C<br>S N G I P Y **A** H W Y Q |
| VLCDR1RHS_H176A | AGC AAC GGT ATT CCG TAC CTG gcC TGG TAT CAG C<br>S N G I P Y L **A** W Y Q |
| VLCDR1RHS_Y174X | AGC AAC GGT ATT CCG NMT CTG CAC TGG TAT CAG C<br>S N G I P **X** L H W Y Q |
| Additional oligos for gap-filling: | |
| VLCDR1RHS_Y174HP | AGC AAC GGT ATT CCG cmT CTG CAC TGG TAT CAG C |
| VLCDR1RHS_Y174ST | AGC AAC GGT ATT CCG wcT CTG CAC TGG TAT CAG C |

| Name | Sequence |
|---|---|
| **VL CDR2** | |
| VLCDR2_r (new) | GATCAGCAGCTTCGGCGCTTTG, Tm=67°C, Ta=70°C |
| VLCDR2_Parent f | TAC CGT GTG AGC AAC CGT TTC AGC GGT GTT CCG A<br>Y R V S N R F S G V P |
| VLCDR2_Y191A | gcC CGT GTG AGC AAC CGT TTC AGC GGT GTT CCG A<br>**A** R V S N R F S G V P |
| VLCDR2_R192A | TAC gcT GTG AGC AAC CGT TTC AGC GGT GTT CCG A<br>Y **A** V S N R F S G V P |
| VLCDR2_V193A | TAC CGT GcG AGC AAC CGT TTC AGC GGT GTT CCG A<br>Y R **A** S N R F S G V P |
| VLCDR2_S194A | TAC CGT GTG gcC AAC CGT TTC AGC GGT GTT CCG A<br>Y R V **A** N R F S G V P |
| VLCDR2_N195A | TAC CGT GTG AGC gcC CGT TTC AGC GGT GTT CCG A<br>Y R V S **A** R F S G V P |
| VLCDR2_R196A | TAC CGT GTG AGC AAC gcT TTC AGC GGT GTT CCG A<br>Y R V S N **A** F S G V P |
| VLCDR2_F197A | TAC CGT GTG AGC AAC CGT gcC AGC GGT GTT CCG A<br>Y R V S N R **A** S G V P |
| VLCDR2_S198A | TAC CGT GTG AGC AAC CGT TTC gcC GGT GTT CCG A<br>Y R V S N R F **A** G V P |
| VLCDR2_Y191X | NMT CGT GTG AGC AAC CGT TTC AGC GGT GTT CCG A<br>**X** R V S N R F S G V P |
| VLCDR2_R192X | TAC NMT GTG AGC AAC CGT TTC AGC GGT GTT CCG A<br>Y **X** V S N R F S G V P |
| VLCDR2_N195X | TAC CGT GTG AGC NMT CGT TTC AGC GGT GTT CCG A<br>Y R V S **X** R F S G V P |
| Additional oligos for gap-filling: | |
| VLCDR2_Y191DHN | vaT CGT GTG AGC AAC CGT TTC AGC GGT GTT CCG A |
| VLCDR2_R192A | TAC gcT GTG AGC AAC CGT TTC AGC GGT GTT CCG A |
| VLCDR2_R192H | TAC caT GTG AGC AAC CGT TTC AGC GGT GTT CCG A |
| VLCDR2_N195PT | TAC CGT GTG AGC mcT CGT TTC AGC GGT GTT CCG A |
| VLCDR2_N195Y | TAC CGT GTG AGC taT CGT TTC AGC GGT GTT CCG A |

| Name | Sequence |
|---|---|
| **VL CDR3** | |
| VLCDR3_f | GCAATAGTAGGTCGCAAAATCC,6 Tm=4°C, Ta=65°C |
| VLCDR3_Parent f | AGC CAG GGT ACC CAT GTG CCG CCG ACC TTC GGT CAA G<br>S Q G T H V P P T F G Q |
| VLCDR3_S231A | gcC CAG GGT ACC CAT GTG CCG CCG ACC TTC GGT CAA G<br>**A** Q G T H V P P T F G Q |
| VLCDR3_Q232A | AGC gcG GGT ACC CAT GTG CCG CCG ACC TTC GGT CAA G<br>S **A** G T H V P P T F G Q |
| VLCDR3_G233A | AGC CAG GcT ACC CAT GTG CCG CCG ACC TTC GGT CAA G<br>S Q **A** T H V P P T F G Q |
| VLCDR3_T234A | AGC CAG GGT gcC CAT GTG CCG CCG ACC TTC GGT CAA G<br>S Q G **A** H V P P T F G Q |
| VLCDR3_H235A | AGC CAG GGT ACC gcT GTG CCG CCG ACC TTC GGT CAA G<br>S Q G T **A** V P P T F G Q |
| VLCDR3_V236A | AGC CAG GGT ACC CAT GcG CCG CCG ACC TTC GGT CAA G<br>S Q G T H **A** P P T F G Q |
| VLCDR3_P237A | AGC CAG GGT ACC CAT GTG gcG CCG ACC TTC GGT CAA G<br>S Q G T H V **A** P T F G Q |
| VLCDR3_P238A | AGC CAG GGT ACC CAT GTG CCG gcG ACC TTC GGT CAA G<br>S Q G T H V P **A** T F G Q |
| VLCDR3_T239A | AGC CAG GGT ACC CAT GTG CCG CCG gcC TTC GGT CAA G<br>S Q G T H V P P **A** F G Q |
| VLCDR3_G233X | AGC CAG NMT ACC CAT GTG CCG CCG ACC TTC GGT CAA G<br>S Q **X** T H V P P T F G Q |
| VLCDR3_P238X | AGC CAG GGT ACC CAT GTG CCG NMT ACC TTC GGT CAA G<br>S Q G T H V P **X** T F G Q |
| Additional oligos for gap-filling: | |
| VLCDR3_G233T | AGC CAG acT ACC CAT GTG CCG CCG ACC TTC GGT CAA G |
| VLCDR3_P238S | AGC CAG GGT ACC CAT GTG CCG tcT ACC TTC GGT CAA G |
